# *In situ* protein crystallography: a single-crystal electron diffraction pipeline for structure determination inside living cells

**DOI:** 10.1101/2025.07.08.663504

**Authors:** Štěpánka Bílá, Dominik Pinkas, Krishna Khakurel, Juliane Boger, Tomáš Bílý, Janos Hajdu, Zdeněk Franta, Roman Tůma, Lars Redecke, Vitaly Polovinkin

## Abstract

Intracellular crystallization is an emerging approach in structural biology that bypasses the need for protein purification. An *InCellCryst* pipeline was recently established for structure determination by serial X-ray crystallography. Serial crystallography requires the exposure of tens of thousands of cells containing intracellular crystals, precluding high-resolution structural studies on proteins with low numbers of crystals. To overcome this limitation, we combined the *InCellCryst* approach with the advantages of 3D electron diffraction and established the new *IncellED* method. Using microcrystals of the HEX-1 protein from *Magnaporthe grisea*, grown inside High Five insect cells, we demonstrate that electron diffraction data collected from a single crystal yields a high-resolution structure at 1.9 Å resolution, comparable to 1.8 Å achieved by serial X-ray crystallography. The *IncellED* methodology opens structure determination for proteins that crystallize with low efficiency and use widely available electron cryo-microscope infrastructure, paving the way for intracellular macromolecular electron crystallography at high resolution.

## Introduction

Protein crystallization within living cells occurs naturally, mainly being associated with various cellular functions such as storage, protection, solid-state catalysis, and sometimes result from progression of diseases and pathogenic infections^1^. It has been demonstrated that intracellular crystallization can be induced by recombinant gene expression in host cells. This emerging approach of structural biology, also referred to as *in cellulo* crystallization^1,2^, complements the diversity of conventional methods of protein crystallization^1,2^. The quasi-native environment within the cell enables the binding and unique identification of physiologically relevant ligands and cofactors of the enzyme^2,3^, which currently cannot be replaced by AI-based structure prediction algorithms. Moreover, crystallization of fully glycosylated proteins, *e.g. T. brucei* cathepsin B, have been observed in living insect cells^4,5^, while conventional crystallization trials were not successful so far. Since the first structure determined in 2007 for a recombinantly produced and intracellularly crystallized protein^6^, the overall approach for structure elucidation of *in cellulo* crystallized recombinant proteins has evolved significantly. Initially, the small width (<10 μm) of the frequently formed needle-shaped crystals posed a significant challenge for X-ray crystallography, since the irradiated crystal volume did not diffract at high resolution using conventional X-ray sources. This has been addressed by the development of micro - and nano-focus beamlines at third- and fourth-generation synchrotron facilities, and the commissioning of X-ray free-electron lasers (XFELs). Combined with novel serial data collection strategies, these technical advances enabled studying of these tiny intracellular crystals at the atomic level^7^, resulting in the deposition of more than 30 high-resolution structures obtained by this approach in the PDB^3,5,6,8–24^.

Streamlined approaches for systematic testing of target protein crystallization within live insect cells have been proposed^14,25^, including the recently published *InCellCryst* pipeline^2^. A recombinant baculovirus is employed to drive target gene expression to increase local protein concentrations, triggering the crystal nucleation process and efficient crystal growth^2,4,5,26,27^. The structure is then determined via serial X-ray diffraction of the micro-crystals directly in the living cells^2^. This approach preserves the native state of the biomolecules and bypasses the need for laborious target protein purification. However, a significant limitation of the intracellular crystallization approaches is the crystallization efficiency, i.e. percentage of cells containing at least one target protein crystal in the infected culture. Testing of novel target proteins frequently resulted in the formation of ordered crystal-like structures, but only in a few cells within the entire culture, sometimes in less than 0.1%. Since the small size of the intracellular crystals requires single exposures of thousands of individual crystals to collect a full diffraction dataset, the low number of crystal-containing cells often precludes structure elucidation for many target proteins even at advanced synchrotron and XFEL sources, constituting the major bottleneck of the *InCellCryst* approach.

Benefiting from strong interactions of electrons with matter^28^, 3D electron diffraction (3D ED; also referred to as MicroED)^29,30^ allows determination of protein structures from sub-micrometer-sized 3D crystals (*e.g.*, a lysozyme structure at 2.1 Å resolution^31^ was obtained from a single crystal with a volume of just 0.14 µm³), which is not possible with other structural biology techniques. Advancements in 3D ED have yielded a number of new protein and peptide structures (*e.g.* see^29,32–36^) and have led to the technical ability to achieve atomic resolution (better than 1.2 Å^7^) for protein structures just from ED data collected from a single crystal with sub-micrometer thickness or a few of such crystals combined^37,38^. The very small size and low number of crystals required makes 3D ED a promising technique for structure elucidation of *in cellulo* crystallized proteins, overcoming the major bottleneck imposed by the low crystallization efficiency and small crystal size.

In a typical 3D ED/MicroED experiment^29,39^, a protein crystal grown by conventional *in vitro* techniques is deposited onto an transmission electron microscopy (TEM) grid, vitrified, and an ED diffraction dataset is recorded under cryogenic conditions (cryo-TEM) as the crystal is continuously rotated under parallel illumination with a high-energy electron beam (typically, 200 or 300 kV). The optimal crystal thickness for the 3D ED collection is in the range of 200-300 nm^40,41^. This poses a problem for thicker samples (*e.g.* crystals within cells), for which the ED signal is strongly affected by multiple scattering and absorption^29^. Thus, thinning larger crystals or cellular samples to lamellae of desired thickness is required and can be performed by cryogenic focused ion beam milling (cryo-FIB milling) within dual-beam focused ion beam scanning electron (FIB/SEM) microscopes^42–44^. Diverse protein crystals were demonstrated to retain a high degree of crystalline order after FIB thinning^37,45,46^, making cryo-FIB milling a standard and reliable method for MicroED sample preparation.

Cryo-FIB/SEM microscope, which only provides topological information about the sample, is not suitable for localizing the target *in cellulo* crystals that are often buried deep within the cell bulk. In principle, this problem can be solved by adapting cryo-light microscopy (cryo-LM) and correlative light-electron microscopy (CLEM) techniques that were previously developed for localizing regions of interest for cellular cryo-electron tomography (cryo-ET)^44,47^.

In the present work, we have built on the aforementioned concepts and extended the *InCellCryst* pipeline to allow 3D ED data collection from single crystals, establishing the new “ *IncellED*” method. Using microcrystals of the HEX-1 protein from the filamentous fungus *Magnaporthe grisea* (*Mg*HEX-1) grown inside High Five insect cells, we demonstrate that our approach delivers cellular lamellae containing portions of the target crystals, suitable for high-resolution structure determination by 3D ED. ED data from ∼1.6 µm³ and ∼0.8 µm³ volumes yielded the previously unknown structure of *Mg*HEX-1 at resolutions of 1.9 Å and 2.2 Å, respectively. These structures were compared to an X-ray structure obtained at 1.8 Å resolution by serial synchrotron-radiation diffraction (SSX) using tens of thousands of *Mg*HEX-1 crystals in High Five cells. The *IncellED* method elucidated similar structural details just from a single crystal. This opens the intracellular crystallization approach to proteins that crystallize with low efficiency and thus overcomes the most significant bottleneck of the *InCellCryst* approach.

## Results

### Intracellular crystallization of *Mg*HEX-1

To establish the *IncellED* approach, *Mg*HEX-1, a structurally uncharacterized HEX-1 protein derived from the filamentous fungus *Magnaporthe grisea*, has been selected. HEX-1 proteins are highly conserved among ascomycetes and self-assemble into a hexagonal crystalline core within peroxisome -derived organelles (Woronin bodies) and help to seal septal pores during cell stress^48–50^. A *Neurospora crassa* HEX-1 (*Nc*HEX-1) was found to produce regular, micrometre-sized hexagonal crystals when overexpressed in insect cells using the recombinant baculovirus (rBV) system^51^ and was quickly established as a test protein for intracellular crystal detection by various techniques (*e.g.*, small angle X-ray scattering - X-ray powder diffraction, SAXS-XRPD^51^, fixed-target serial diffraction data collection at XFELs^16^ and synchrotron sources^52^) and led to the development of the *InCellCryst* pipeline^2^. The crystallization efficiency and low-micrometre size range position HEX-1 crystals as ideal candidates for *IncellED* method development. Moreover, the elucidation of a second HEX-1 structure can provide new insights into biological self -assembly processes and into the structural conservation within this protein family. Thus, we selected *Mg*HEX-1 as a pilot system in this study that shares 74% sequence identity with *Nc*HEX-1.

Applying the *InCellCryst* approach, up to three hexagonal-shaped structures per cell were detected by light microscopy in approximately 40% of the rBV-infected High Five insect cells at day four post infection, exhibiting a hexagonal morphology similar to that of previously reported *Nc*HEX-1 crystals^2,50^ (**Supplementary Fig. 1**). The mean size of *Mg*HEX-1 crystals, which ranged from 4 to 15 µm in the longest dimension, was estimated at 8.6 ± 2.5 µm (± standard deviation) (**Fig. 1a, b**). Infected High Five cells also produce EYFP (enhanced yellow fluorescent protein) as a fluorescence marker encoded on the EmBacY bacmid^53^, which enables 3D visualization of intracellular *Mg*HEX-1 crystals in a negative imaging manner using fluorescence (FL) microscopy (**Fig. 1c, d, e**). This was used for 3D targeting of crystals in subsequent cryo-FIB experiments. The High Five cells containing the *Mg*HEX-1 crystals were used both for 3D ED experiments and for X-ray serial crystallography measurements.

**Fig. 1.**
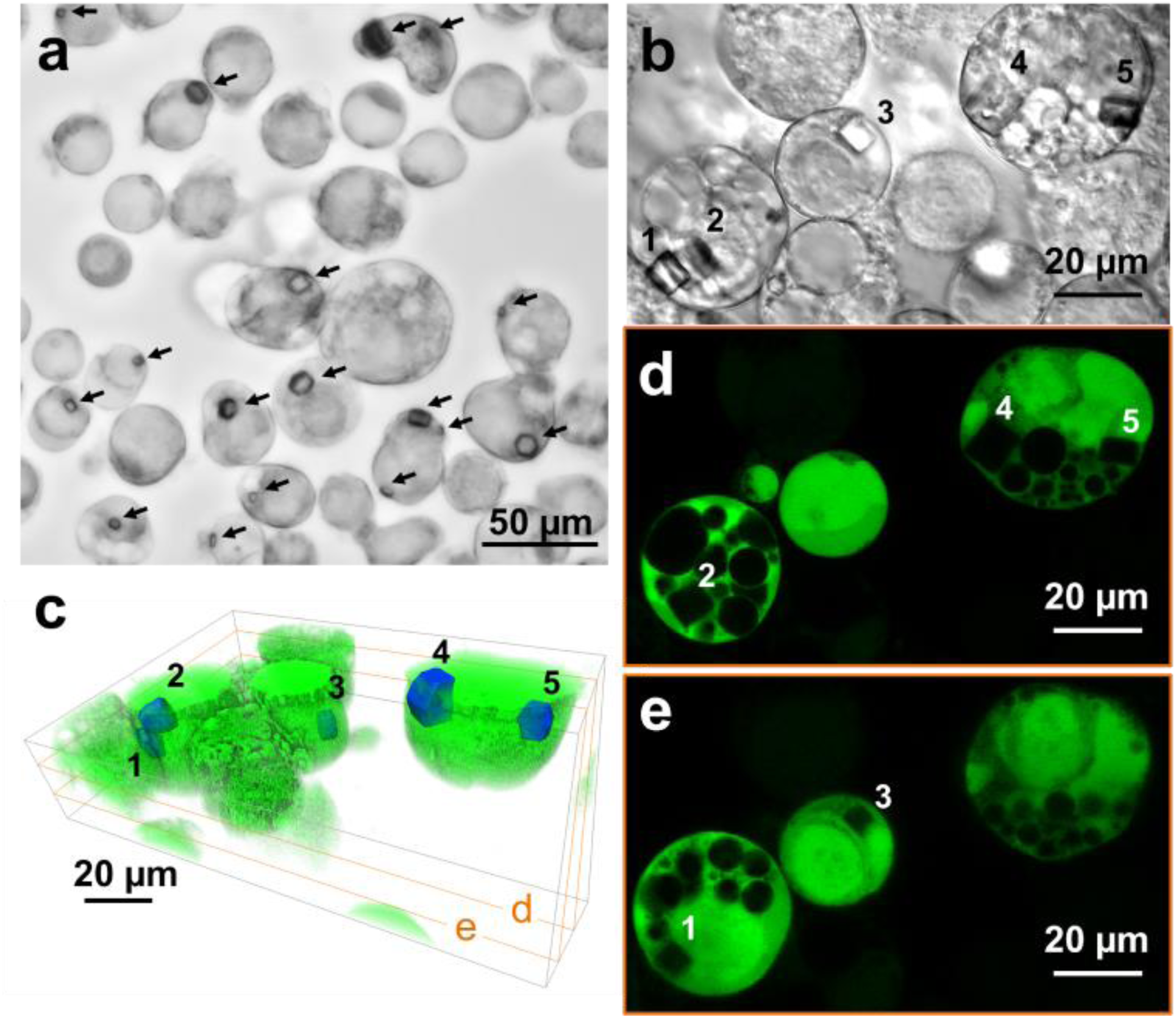
Intracellular crystallization of *Mg*HEX-1 in High Five insect cells and negative fluorescence (FL) imaging of the *in cellulo* crystals. (**a**) A median intensity Z-projection of bright-field imaging Z-stacks shows the cell suspension used for ED and serial X-ray crystallography experiments. Hexagonal-shaped *Mg*HEX-1 crystals are indicated by black arrows. (**b**) A median intensity Z-projection of a bright-field Z-stack of a smaller region of the cell suspension, with numbers indicating individual crystals. (**c**) 3D model visualization of the region in (**b**) based on a confocal FL microscopy imaging Z-stack acquired using 488 nm laser excitation at room temperature. The FL excitation of enhanced yellow fluorescent protein (EYFP), co -expressed with *Mg*HEX-1, allows for the detection of the *in cellulo* crystals as “sharp-edged” areas lacking the FL signal. The crystals are blue-colored for enhanced visualization. The FL virtual Z-slices in (**d**) and (**e**) show areas corresponding to the *Mg*HEX-1 crystals labelled by numbers in (**b**) and lacking the EYFP FL signal.

### *In cellulo* crystal preparation for ED experiments

Building upon reported crystal cryo-FIB milling procedures^44,45,54^ and the idea of using 3D negative FL imaging for accurate 3D localization of target crystals within a cryo-FIB/SEM microscope, we established the protocol of ED sample preparation. The procedure is shown schematically in **Fig. 2** and described in detail in *Methods*.

**Fig. 2.**
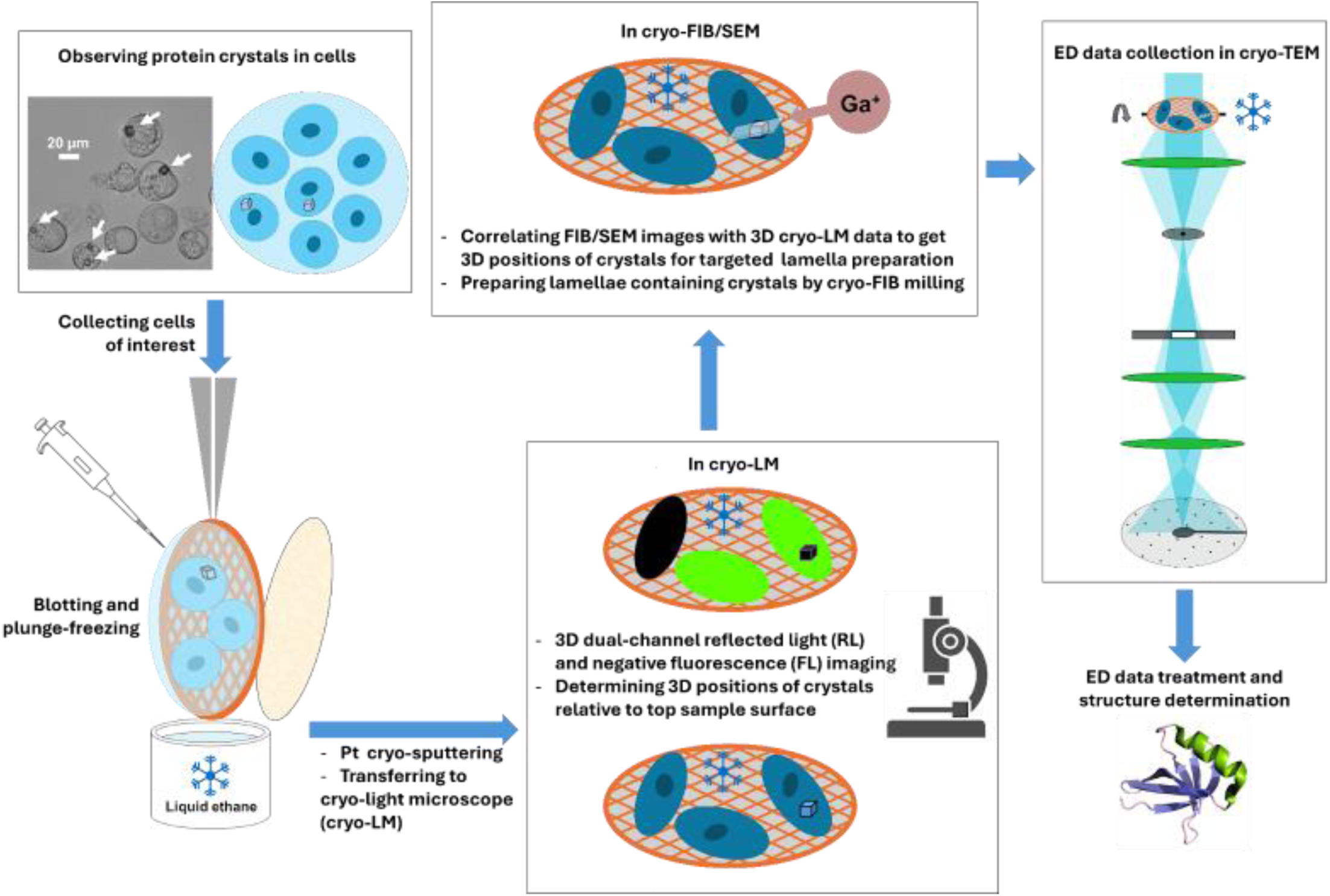
Schematic diagram of the *IncellED* experimental workflow.

Presence of *Mg*HEX-1 crystals in cells was verified by light microscopy prior to TEM grid preparation and plunge-freezing. The frozen cells on the grid underwent 3D imaging by a cryo -light microscope (cryo-LM) (**Supplementary Fig. 2 and 3**). Negative FL imaging yielded 3D coordinates of crystal centers relative to the sample’s top surface, established by reflected light (RL) imaging (**Supplementary Fig. 3**). The 3D RL signal provided surface topography analogous to FIB/SEM imaging. Both RL contrast and subsequen t FIB/SEM imaging were improved by metallic Pt sputter coating (see *Methods* for details). Within the cryo-FIB/SEM microscope, the RL data with the 3D positions of the crystals, referenced to the sample surface, were correlated with FIB/SEM images via shared surface features (**Supplementary Fig. 4 and 5**). This enabled accurate localization of target crystals for subsequent cryo-FIB milling. The selected *Mg*HEX-1 crystals were typically in the range of 5-8 µm (see *e.g.* **Fig. 3a**), slightly below the average observed crystal size of 8.6 µm, and only the crystals fully contained within the cell were used for subsequent processing. Lamellae containing the parts of target crystals were prepared by cryo-FIB milling with 30 kV Ga^+^ ions (**Fig. 3b, c**; **Supplementary Fig. 6 and 7**) and used for 3D ED measurements.

**Fig. 3.**
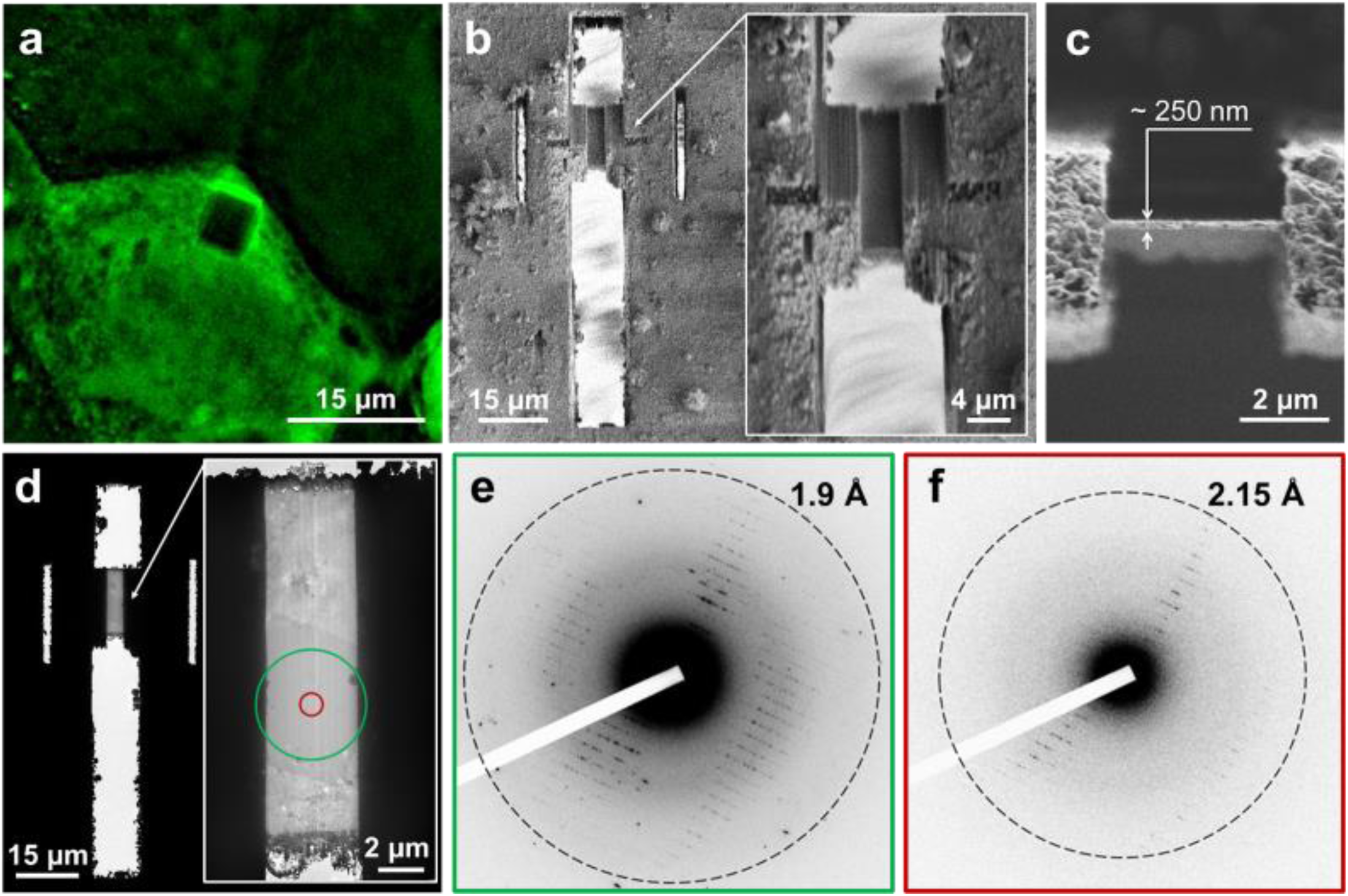
Example of a prepared lamella containing an *in celullo Mg*HEX-1 crystal and its ED characterization in a 200 kV cryo-TEM. (**a**) EYFP-derived negative FL image of the targeted *Mg*HEX-1 crystal, acquired around its Z-center under cryogenic conditions before the Ga^+^ FIB-milling procedure. For ED analysis, the lamella was machined around the crystal center, had a thickness of ∼250 nm, a width of ∼4 µm, and was angled at 20° to the TEM grid plane. (**b**) 0.5 kV SEM image of the lamella prepared by cryo-FIB-milling; the inset shows an SEM image of the lamella with higher resolution. **(c)** 30 kV Ga^+^ FIB image of the lamella. (**d**) TEM image of the FIB-milled lamella containing a portion of the targeted *Mg*HEX-1 crystal; the inset shows a TEM image of the lamella with higher resolution. (**e**) A 200 kV electron diffraction (ED) pattern obtained from a ∼4.8 µm area of the lamella, defined by the parallel beam illumination and depicted by a green circle in ( **d**). The crystal volume impacting the (**e**) ED signal was estimated to be ∼4.4 µm³ when excluding non-electron- transparent (black) parts in (**d**). (**f**) A 200 kV ED pattern collected from a 1 µm area of the crystal (crystal volume ∼0.2 µm³), defined by a selected area aperture of the cryo -TEM and depicted by a red circle in (**d**). Both the ED signals were acquired at the same electron beam fluence of ∼0.08 e/Å².

### Electron diffraction data collection

First, a lamella containing a fragment of an *in cellulo* grown *Mg*HEX-1 crystal (∼6 µm in size; **Fig. 3**) was used to assess whether the sample preparation method yields ED data suitable for high -resolution structure determination. The lamella was prepared with a thickness of 250 nm, an optimal value previously reported for 200 kV ED^40,41^. Following localization of the crystal in imaging mode within a 200 kV cryo -TEM (**Fig. 3d**), ED data were recorded as still images using a scintillator-based electron detector (see *Methods*). At the same electron fluence of ∼0.08 e/Å², high-resolution ED signals up to 1.9 Å (**Fig. 3e**) and 2.15 Å (**Fig. 3f**) resolutions were detected from a ∼4.8 µm diameter zone (**Fig. 3d** green circle, crystal volume of ∼4.4 µm^3^) and a 1 µm diameter zone (**Fig. 3e, f** red circle, crystal volume of ∼0.2 µm^3^), respectively. Thus, the sample preparation method yields crystal lamellae diffracting to high-resolution even at low crystal volumes while using scintillator-based (indirect) electron detection. However, direct electron detection (DED) combined with energy filtration provides superior sensitivity and signal-to-noise ratios, yielding a significantly improved ED data^38,55^. Thus, data for *Mg*HEX-1 structure determination was collected using a 300 kV microscope equipped with DED K3 electron-counting camera in a continuous rotation mode while filtering out inelastically scattered electrons with energy loss more than 10 eV (see *Methods* for full detail).

We produced four lamellae containing crystals ranging from 5 to 8 µm in size for the 3D ED structure determination experiments. The thickness of the four cryo-FIB milled lamellae was ∼300 nm, i.e. optimal for 300 kV ED^40,41^. For two of the lamellae, the 3D ED data were acquired using a 2.5 µm diameter collection zone (volume of ∼1.6 µm^3^; denoted as “microvolume”) with a total fluence of ∼0.3 e/Å². This corresponds to ∼1.1 MGy dose absorbed, which is much below the doses reported for observing radiation dama ge using ED^56,57^. These microvolume 3D ED datasets were characterized by highly similar data processing statistics, and one of them was selected for *Mg*HEX-1 structure elucidation via molecular replacement (MR). Previously reported *Nc*HEX-1 structure (PDB 1KHI)^50^, which shares 74% sequence identity, was used as a search model for MR. This yielded a 1.9 Å resolution crystal structure of *Mg*HEX-1 with P6_5_22 symmetry (**Table 1** and **Fig. 4a**), as previously reported for *Nc*HEX-1^50^. The structure model was refined to 0.205/0.229 of R_work_/R_free_.

**Fig. 4.**
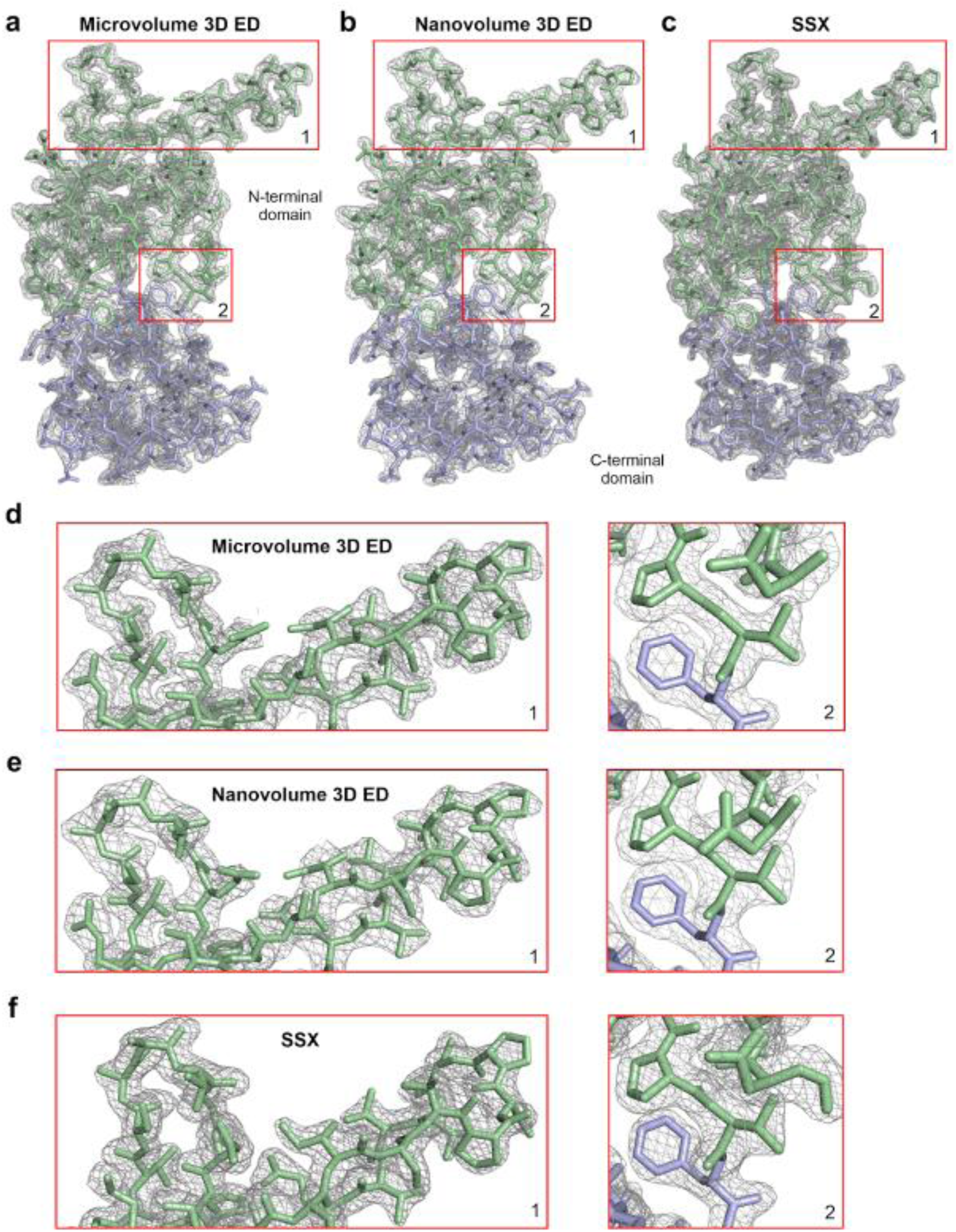
High-quality electrostatic potential and electron density maps obtained from micro - and nanovolume 3D ED and serial synchrotron-radiation X-ray crystallography (SSX) data collected from *in cellulo Mg*HEX-1 crystals. Electrostatic scattering potential maps 2Fo-Fc contoured at 1*σ* (grey) calculated from the micro- (**a**) and nanovolume (**b**) 3D ED data. The modelled *Mg*HEX-1 protein structures are shown in green (N-terminal domain) and blue (C-terminal domain) stick representation. For direct comparison, the electron density map obtained from serial X-ray diffraction of approximately 62,500 intracellular *Mg*HEX-1 crystals is presented (**c**). Selected regions (1, residues Thr65 to His72 and Ser88 to Gln101; 2, residues Gly105 to Lys109) of the electrostatic potential and electron density maps (red squares) are shown in detail for the micro - (**d**) and the nanovolume 3D ED data (**e**) as well as for the SSX data (**f**).

**Table 1.** X-ray and 3D ED/MicroED data collection, processing and refinement statistics for *Mg*HEX-1 crystal structures.

| **Data set** | **Serial synchrotron X-ray diffraction (SSX)** | **3D ED microvolume** | **3D ED nanovolume** |
| --- | --- | --- | --- |
| **Number of crystals** | 62,496 | 1 | 1 |
| **Dose per crystal (MGy)** | ~1.4 | ~1.1 | ~2.2 |
| **Sample temperature (K)** | 100 | 80 | 80 |
| **Processing software** | CrystFEL | XDS, AIMLESS | XDS, AIMLESS |
| **Space group** | P6<sub>5</sub>22 | P6<sub>5</sub>22 | P6<sub>5</sub>22 |
| **Unit cell (Å or °)** | 57.66 57.66 187.89<br>90 90 120 | 57.83 57.83 198.18<br>90 90 120 | 57.38 57.38 198.12<br>90 90 120 |
| **Resolution range (Å)** | 93.95 - 1.80<br>(1.82 - 1.80) | 11.94 - 1.90<br>(1.95 - 1.90) | 11.68 - 2.20<br>(2.27 - 2.20) |
| **Total reflections** | 31,949,243 | 123,226 | 43,560 |
| **Unique reflections** | 18,144 (855) | 16,281(1,053) | 9,305 (835) |
| **Multiplicity** | 176.9 (57.2) | 7.6 (8.0) | 4.7 (5.0) |
| **Completeness (%)** | 100.00 (99.88) | 99.4 (98.5) | 91.7 (92.8) |
| **I/σI** | 25.86 (0.59) | 4.8 (0.4) | 3.7 (0.5) |
| **R-split (%)** | 3.34 (177.35) | - | - |
| **R-pim (%)** | - | 11.3 (151.6) | 13.1 (103.3) |
| **CC1/2** | 0.9993 (0.2583) | 0.989 (0.475) | 0.973 (0.655) |
| **CC\*** | 0.9998 (0.6407) | - | - |
| **No. collected images** | 110,030 | - | - |
| **No. hits / indexed lattices** | 59,042/62,496 | - | - |
| **Refl. used in Refinement** | 17,256 (1,322) | 15,389 (1,076) | 8,745 (603) |
| **Refl. used for R-free** | 1,401 (108) | 809 (55) | 499 (38) |
| **R-work** | 0.2130 (0.4174) | 0.2052 (0.3170) | 0.2111 (0.3660) |
| **R-free** | 0.2364 (0.4764) | 0.2292 (0.3650) | 0.2576 (0.4300) |
| **No. non-hydrogen atoms** | 1,126 | 1190 | 1,171 |
| **Macromolecules** | 1,033 | 1,124 | 1,101 |
| **Solvent** | 93 | 66 | 70 |
| **RMS bonds (Å)** | 0.009 | 0.008 | 0.005 |
| **RMS angles (°)** | 1.05 | 1.57 | 1.30 |
| **Ramachandran favored (%)** | 98.52 | 97.90 | 95.10 |
| **Ramachandran allowed (%)** | 1.48 | 2.10 | 4.90 |
| **Ramachandran outliers (%)** | 0 | 0 | 0 |
| **Rotamer outliers (%)** | 1.77 | 0.80 | 0 |
| **Clash score** | 6.81 | 4.43 | 3.18 |
| **Average B-factor (Å<sup>2</sup>)** | 40.54 | 32.10 | 51.00 |
| **Macromolecules** | 40.05 | 31.68 | 52.06 |
| **Solvent** | 45.92 | 33.64 | 48.62 |
| **Mean RMSD to 1KHI (Å)** | 0.520<br>(134 atoms) | 0.341<br>(134 atoms) | 0.340<br>(134 atoms) |

The other two lamellae were tested using a 1.75 µm in diameter (volume of ∼0.8 µm^3^; denoted as “nanovolume”) collection zone. The 3D ED data were collected from one lamella at a total fluence of ∼0.6 e/Å². This corresponds to ∼2.2 MGy radiation dose absorbed, again below the dose limit for observing radiation damage using ED^56,57^. Processing of the nanovolume single crystal ED data resulted in a 2.2 Å resolution structure of *Mg*HEX-1 that was refined to 0.211/0.258 of R_work_/R_free_ (**Table 1** and **Fig. 4b**). Interestingly, the nanovolume ED signal from the second lamella was observed up to 1.7 Å and exhibited the highest resolution detected for all lamellae tested in this study at the same electron fluence (**Supplementary Fig. 8**). Unfortunately, this lamella was contaminated by ice microcrystals (**Fig. 8c**) during cryo-transfer to the cryo-TEM from storage, preventing reasonable 3D ED data collection for structure determination. Considering the variation in diffraction strength among the four examined *in cellulo* crystal lamellae, analyzing additional crystals presents a promising opportunity to achieve higher resolution and quality of 3D ED structure. This potential remains to be explored in future studies.

### 3D ED structure of MgHEX-1

The electrostatic potential maps were refined for the 3D ED micro - and nano-volume datasets at 1.9 Å and 2.2 Å resolution (**Fig. 4a, b** and **Table 1**) and provided a high level of detail, sufficient to rebuild the non-flexible part of the main *Mg*HEX-1 chain and to model most of the side chains. No interpretable electrostatic potential density is observed for the N-terminal residues 1–35, which have already been classified as disordered for the *Nc*HEX-1 structure^50^, and the three C-terminal residues 179-181. The maps of Gly153, Arg154 and Gly155 that are located in a flexible loop at the C-terminus of helix 1 are not fully resolved, as well as the side chains of residues Gln117 and Arg148. The final micro - and nano-volume models show a protein backbone Cα root mean square deviation (RMSD) of 0.341 Å and 0.340 Å, respectively, compared to the 134 equivalent Cα atoms of the MR search model, which is in the expected range for proteins of this sequence identity level. Moreover, the micro- and nano-volume ED models are almost superimposable with an overall RMSD of 0.166 Å for the 123 Cα atoms. Deviations of more than 0.5 Å are restricted to residues Glu151 and Ser152 located in the flexible loop mentioned above. Even for the side chains the RMSD does not exceed 0.75 Å, except for the three residues Asn133, Glu144, Arg148 that are not fully resolved in both maps.

### X-ray crystal structure of *Mg*HEX-1 and comparison to 3D ED structures

Next, we compared the structural information obtained by 3D ED from a single micro - or nano-volume *Mg*HEX-1 crystals to that obtained by applying the well-established approach of serial synchrotron X-ray diffraction (SSX) on the crystal-containing insect cells^2^. Visual inspection of test X-ray patterns revealed diffraction up to 2 Å, allowing the collection of a complete, serial line scan dataset. Data processing using CrystFEL and MR using again the *Nc*HEX-1 structure (PDB 1KHI) as a search model yielded an electron density map at 1.8 Å resolution, which allowed refinement of the structure model to 0.213/0.236 of R_work_/R_free_ (**Table 1**, **Fig. 4c**). The final *Mg*HEX-1 structure included residues 35 to 174, again without the flexible N-terminus. In contrast to the 3D ED maps, no interpretable electron density is observed for residues 153 and 154 located in the previously mentioned flexible loop, and for the C-terminal residues 175 to 178. Likewise, the side chains of residues Arg112 and Gln117 are not fully resolved. The calculated RMSD of 0.520 Å compared to the 134 equivalent Cα atoms of the MR search model is increased compared to that of the 3D ED structures, but still in the expected range.

Comparison of the 3D ED micro- (RMSD 0.546 Å for 134 equivalent Cɑ atoms) and nano-volume *Mg*HEX-1 structures (RMSD 0.509 Å for 134 equivalent Cɑ atoms) obtained from single crystals with that elucidated by SSX using 62,496 intracellular crystals revealed no major differences in the overall conformation of the *Mg*HEX-1 monomer. Deviations of morethan 1 Å are restricted to residues Ser66 to Gly70 located in a flexible loop connecting *β*-strands 3 and 4 in the N-terminal domain (**Fig. 5a**). Also, the calculated overall RMSD for the side chain atom positions compared to the micro-volume (0.738 Å) and the nano-volume (0.756 Å) model indicated an almost identical *Mg*HEX-1 structure obtained by 3D ED and SSX.

**Fig. 5.**
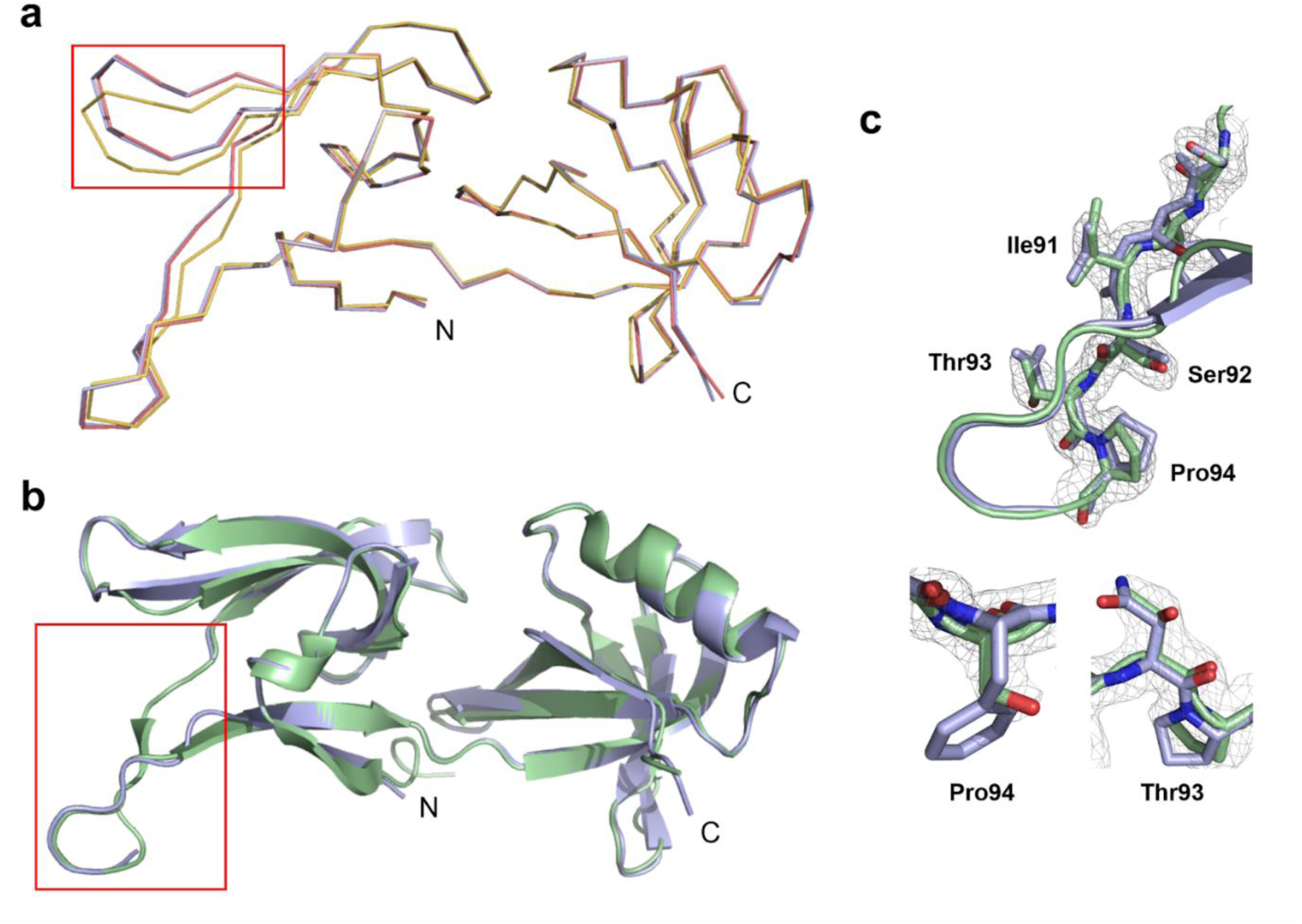
Structural homology of HEX-1 proteins. (**a**) Superposition of the *Mg*HEX-1 3D ED micro- (blue) and nano-volume (red) structures with the *Mg*HEX-1 SSX structure (yellow), all in backbone representation. The only region showing major structural differences with RMSDs above 1.0 Å is highlighted by the red box. ( **b**) Cartoon representation of the *Nc*HEX-1 reference structure (PDB 1KHI; green) purified from *E. coli* and crystallized by sitting-drop vapour diffusion superimposed with that of *Mg*HEX-1 crystallized in insect cells and obtained from microvolume 3D ED (blue). The average RMSD is 0.45 Å for 123 equivalent C atoms. (**c**) 2Fo-Fc map (grey, contoured at 1*σ*) of most deviant residues 89-94 (red box in (**b**)) of the *Mg*HEX-1 3D ED micro-volume structure (green) in stick representation superimposed with the corresponding residues in the *Nc*HEX-1 structure (blue). The side chains of residues Ser90 and Thr93 fit well into the 2Fo-Fc map calculated for *Mg*HEX-1, in contrast to the corresponding Phe and Asn residues of *Nc*HEX-1, ruling out phase bias imposed by phase determination applying the MR approach with a similar reference structure as the search model.

The observed difference in the c-axes of the unit cells between the SSX structure and the 3D ED structures (**Table 1**) is likely attributable to the impact of distinct sample handling conditions. For SSX, cells were overlaid with PEG200 for cryo-protection during freezing in liquid nitrogen, which causes dehydration. For 3D ED experiments, cells were plunge-frozen in liquid ethane, which does not require cryo-protection due to a faster freezing process. A similar effect was previously observed for serial femtosecond X-ray data collection at an XFEL at room temperature^16^.

All *Mg*HEX-1 models revealed the typical two-domain structure that was previously reported for *Nc*HEX-1^50^, consisting of mutually perpendicular antiparallel *β*-barrels (**Fig. 5b**). The N-terminal domain is formed by six antiparallel *β*-strands, while the C-terminal domain consists of a five-stranded *β*-barrel and an α-helix. This indicates a high degree of structural conservation among HEX-1 proteins from different Ascomycota, especially in the ‘common region’ comprising the 148 C-terminal residues. However, for residues 89-94, representing the most divergent part between the *Nc*HEX-1 and the *Mg*HEX-1 structure, the side chains fit well into the 2Fo-Fc map of the *Mg*HEX-1 3D ED microvolume structure, in contrast to the corresponding residues of *Nc*HEX-1 (**Fig. 5c**). This rules out model bias imposed by phase determination applying the MR approach with a search model similar to the elucidated *Mg*HEX-1 structure, confirming the correct interpretation of the map.

Three types of intermolecular surface interactions were previously identified that are responsible for the self-assembly of *Nc*HEX-1 into a crystalline state in Woronin bodies^50^, representing the physiological function of HEX-1 proteins in ascomycetes^49^. All involved residues are conserved in terms of sequence and structure between *Mg*HEX-1 and *Nc*HEX-1 (**Supplementary Fig. 9a**). Again, a combination of salt bridges and hydrogen bonds forms the interactions of monomers, previously denoted as group I and group II interactions, which build up a structural HEX-1 polymer that establishes a helical spiral. Each filament is in contact with six identical neighbouring polymers via hydrogen bonds (group III interactions), finally producing the six -fold symmetry of the *Mg*HEX-1 crystals, identical to that reported for *Nc*HEX-1^50^ (**Supplementary Fig. 9b**).

In conclusion, the comparable level of biologically relevant information obtained by the novel *IncellED* approach presented in this study and by well-established SSX validate 3D ED as a suitable technique to obtain *in situ* high-resolution structural information from a single protein crystal in the very low micrometer size range.

## Discussion

In this study, we established and validated *IncellED*, a workflow for high-resolution structure determination by 3D ED using only a single intracellular protein crystal in the micrometer size range.

Electron crystallography has been demonstrated to yield high-resolution protein structures from individual crystals with sizes insufficient for X-ray crystallography (see^29–32,37,38^). In these cases, it is usually the radiation damage that limits the X-ray intensity and irradiation time and precludes collection of a full dataset from a single protein crystal. Thus, for low-micrometer to nano-sized crystals, serial data collection strategies are required, usually combining individual X-ray diffraction stills of tens of thousands of randomly oriented crystals to obtain a complete data set^58,59^. If the crystals are heterogeneous in diffraction capability, merging the diffraction pattern of single crystals can affect the resolution and quality of the dataset. Therefore, the resulting structural model exhibits increased B factors and poorly resolved average conformations of residues located in flexible regions, as observed for the SSX *Mg*HEX-1 structure. The single crystal 3D ED approach presented here allows screening for the best diffracting crystal by probing the individual electron diffraction capabilities of each crystal, without losing structural details by averaging. Considering the total crystalline volume used and the radiation dose deposited, *IncellED* is >10^5^-fold more efficient than the applied SSX approach, while providing a similar level of structural information (**Table 1** and **Supplementary Text 1**). This remarkable difference likely reflects the distinct scattering nature of electrons compared to X-rays (especially their stronger elastic electron scattering and more favourable inelastic *versus* elastic scattering ratio, well-analyzed in^28^), which also enables single-particle cryo-EM imaging.

In general, *IncellED* structure determination may require data from more than one crystal. In this study, the high symmetry (P6_5_22) of the *Mg*HEX-1 crystals enabled 3D ED data collection of high completeness (>90%; **Table 1**) using ∼80° collection wedges from single crystals. However, practical limitations such as increased crystal lamella thickness at high tilts and the proximity of TEM grid bars (**Supplementary Fig. 2**) restrict the usable cryo-TEM’s stage tilt range to less than the current hardware limits (-70° to +70°)^60–63^. For crystals with lower symmetry, this would prevent achieving more than 90% data completeness desired for reliable structure refinement and model building^63^. For example, a P1 symmetry crystal requires a 180° diffraction wedge for 100% completeness, often necessitating data merging from multiple crystals ^37^ – at least from two crystals if oriented properly^64^. Preferred crystal orientation is a common issue in ED experiments where, for example, plate-like crystals align unfavorably with the TEM grid support when deposited onto the grid *in vitro,* thereby creating regions of reciprocal space inaccessible for data collection ^61–63^. However, such a risk is low for *IncellED* since the 3D cellular environment effectively randomizes the crystal orientations (see *e.g.* **Supplementary Fig. 1**).

The sample preparation time, primarily consumed by the Ga^+^ cryo-FIB milling process, was evaluated at ∼2-2.5 h/crystal for the specific instrumental layout used in this work (see *Methods*). Such throughput can be problematic, especially if testing is required for tens of *in cellulo* crystals per session that are, for example, highly variable in terms of diffraction quality or crystal morphology due to differences in intracellular growth conditions^2^. However, utilizing recently introduced, but not yet widespread, plasma FIBs with noble gas ions such as Xe^+^ or Ar^+^, which are more efficient at sputtering cryo-material, and automating the FIB milling process can significantly increase the sample preparation rate, likely reaching a rate of <1 h/crystal and enabling automated overnight runs (see *e.g.*^46,65,66^).

Effective negative fluorescence (FL) imaging of *in cellulo* crystals necessitates the use of a high numerical aperture (NA) objective. This is crucial for resolving small volumes, particularly in the Z-direction, thereby preventing the infiltration of the “positive” FL signal from the surrounding media containing co-expressed EYFP. The current technical implementation of the *IncellED* sample preparation (see *Methods*) employs the negative FL imaging and is expected to provide a crystal center localization precision of ∼1 µm, estimated by considering the Z-axis optical resolution of ∼0.7 µm (the XY- resolution is sub-0.4 µm) and the ∼300 nm Pt-GIS protective coating applied during the sample preparation. This introduces the inaccuracy that prevents sufficient localization precision for *IncellED* preparation of crystals of about 1 µm in size or smaller. However, the *Mg*HEX-1 crystals tested in this study were larger (5-8 µm), and the challenge of preparing sub-1 µm crystals inside cells by targeted cryo-FIB milling was not addressed. This aspect remains to be explored. A potential solution can involve reimaging of the sample lamellae after the “rough”’ milling procedure (resulting in a ∼4.5 µm thick lamella containing a crystal; see *Methods*) with improved 3D optical resolution, for example, by utilizing a confocal cryo-FL microscope^67^ or by preparing cryo-lamellae under the guidance of 3D confocal imaging using a cryo-confocal microscope integrated into the FIB/SEM microscope^68^, thereby achieving a localization precision close to the desired lamella thickness (∼200 -300 nm). As an alternative to employing negative FL imaging, “positive” imaging techniques that directly probe the protein crystal may improve crystal localization precision. Such methods could be based on single- or multi-photon excited UV fluorescence^69–72^, second or third harmonic generation^69,70,73^, and the Raman effect^74,75^, all of which are currently being introduced onto the market. Once implemented in the context of cryo -EM, the positive crystal imaging methods will also adapt the *IncellED* workflow for general usage with crystals naturally occurring within cells that do not possess an inherent or engineered FL signal (like EYFP used in the current setup).

In summary, *IncellED* provides an alternative to the traditional X-ray diffraction methods used for the *in cellulo* protein crystal structural investigation. By requiring several orders of magnitude fewer crystals, *IncellED* uniquely enables the structural analysis of small single crystals. Furthermore, while X-ray diffraction methods often depend on large-scale facilities such as XFELs and synchrotron radiation sources, *IncellED* utilizes a suite of accessible in-house technologies—including cryo-LM, cryo-FIB/SEM, and cryo-TEM with DED. Notably, high-quality 3D ED data can be acquired using widely available 200 kV side-entry TEMs equipped with cryo-sample holders (see *e.g.*^34,76,77^). Given these advantages, we anticipate that the *IncellED* methodology will be broadly adopted within the structural biology community. Last but not least to mention, atomic scattering factors for electrons, unlike those for X-rays, are strongly influenced by the charge state of the scattering atoms^29,76,78^. This makes 3D ED particularly suitable but yet to be explored for mapping charged states of residues and cofactors^79,80^ in proteins crystallized *in cellulo*.

To our knowledge, the present work demonstrates the first high-resolution (sub-2 Å) *in situ* protein structure from a single nanocrystal by electron diffraction. Thus, *IncellED* opens the intracellular crystallization approach to proteins that form small crystals with low efficiency in cells, overcoming the most significant limitation for a more broad application so far. Combined with its potential technical improvements and recent work on investigating the structure of naturally occurring nanocrystals of major basic protein-1 in cells at lower resolution^81^, *IncellED* paves the way for native macromolecular electron crystallography at high resolution using widely available cryo-EM infrastructures.

## Methods

*InCellCryst*, a streamlined approach for *in cellulo* crystallization of recombinant proteins in living insect cells^2^, was applied to *Mg*HEX-1, involving vector construction, virus stock production, virus stock titration, and the detection of protein crystals within insect cells.

### *Mg*HEX-1 vector construction

Cloning of the Hexagonal peroxisome 1 (HEX-1) gene from ascomycetous fungus *Magnaporthe grisea* (NCBI Accession AY044846, codon optimized for *Trichoplusia ni* expression) into the pFastBac1 V2 cyto vector under the control of the polyhedrin promotor^2,82^ was performed by PCR amplification using primers 5’-GATC GGTACCTACTATGAAGACGATCGTGAGACC-3’ and 5’-GATCGCTAGCCAGCCTGCTGCCGTG-3’, adding *Kpn*I and *Nhe*I restriction sites at 5′ and 3′ ends of the *Mg*HEX-1 gene without start and stop codon and HiFi DNA polymerase (highQu). After digestion of the PCR product with FastDigest *Nhe*I and *Kpn*I (Thermo Fisher Scientific), the insert was ligated into the equally digested pFB1 V2 cyto vector by T4 DNA ligase (Thermo Fisher Scientific), yielding pFB1 *Mg*HEX-1 V2 cyto plasmid that additionally encodes EYFP as a marker protein. Competent *E. coli* DH5α cells (NEB C2987) were transformed with ligation mixture and amplified pFB1 *Mg*HEX-1 V2 cyto plasmid DNA was extracted using the GeneJET Plasmid Miniprep Kit (Thermo Fisher Scientific) and sequence verified (LGC Genomics).

### Recombinant baculovirus stock production and titration

For recombinant baculovirus (rBV) production *E. coli* DH10EmBacY cells (Geneva Biotech) were transformed with pFB1 *Mg*HEX-1 V2 cyto according to the Bac-to-Bac manual (Invitrogen). Recombinant bacmid DNA, isolated by applying the ZR Bac DNA Miniprep Kit (Zymo Research), was verified for the presence of the transposed pFB1 sequence by PCR using pUC/M13 primers. In a 12 -well cell culture plate, 5 x 10^5^ *Spodoptera frugiperda* Sf9 insect cells were grown in serum- and protein-free ESF921 cell culture medium (Expression Systems) at 27°C and transfected with recombinant bacmid DNA using Escort IV reagent (Sigma Aldrich). After four days of incubation at 27°C the supernatant was harvested (P1 virus stock) and used for infection of 2 x 10^6^ Sf9 cells/ml in a T-25 ml cell culture flask that was incubated for another 4 days at 27°C with constant shaking (100 rpm), followed by harvesting the supernatant (P2 stock). The virus titre of the P2 stock was calculated using the TCID_50_ (tissue-culture infectious dose^83^) in a serial dilution assay as previously described^2,51,82^. The efficiency of transfection and viral stock production as well as the virus titer was determined by detection of the fluorescence of the EYFP marker encoded in the bacmid^83^ using a widefield inverted X81 (Olympus) or Ts2R-FL (Nikon) microscope.

### *In cellulo* crystallization of *Mg*HEX-1

For intracellular crystallization, 0.9 x 10^6^ High Five insect cells (Thermo Fisher Scientific) were plated in 2 mL ESF921 medium (Expression Systems) in one well of a six-well cell culture plate and infected with the *Mg*HEX-1 encoding rBV using a multiplicity of infection (MOI) of 1. The required amount of P2 stock was calculated using the following formula: virus stock [mL] = (MOI · cell number) / (0,69 · TCID_50_/mL)^84^. After 4 days of incubation at 27°C, the High Five cells were examined for the presence of *Mg*HEX-1 protein crystals by light microscopy via bright field imaging^2^ using Ts2R-FL (Nikon) and DMi8 (Leica Microsystems) inverted microscopes equipped with 20x and 40x objectives, respectively. Median intensity Z-projection of bright-field Z-stacks were performed by ImageJ^85^ to better visualize the crystals. Cells containing *Mg*HEX-1 crystals were carefully harvested by pipetting for subsequent electron or X-ray crystallography experiments. Note that the excitation of EYFP fluorescence (FL), which is co-expressed in the cells, provided an additional way of detecting *in cellulo* crystals as regularly shaped areas lacking the fluorescent signal (**Fig. 1**). At room temperature, the latter was performed with a confocal laser scanning FV1000 microscope (Olympus) using a 488 nm laser and a 100x objective (UPlanSApo, oil-immersion, NA 1.4, Olympus). 3D model visualization based on the FL stack was created using Amira software (Thermo Fisher Scientific). In Amira, the Volume Rendering function and the 2D Median filter were used, and the blue model of the crystals was semi-automatically generated (**Fig. 1**).

### Sample preparation for ED data collection

The sample preparation workflow for 3D ED data collection is depicted in **Fig. 2**. A 3 μl aliquot of cells containing *Mg*HEX-1 crystals was carefully collected with a pipette from the bottom of a well in a six -well cell culture plate. Then a EM GP2 plunge freezer (Leica Microsystems) was used, with its sample environment chamber set to 20°C and 95% relative humidity. The cell aliquot was deposited onto a holey gold film covered TEM grid (UltrAuFoil® R2/2, 200 gold mesh, Quantifoil Micro Tools). The grid was then blotted with filter paper (#1, Whatman) from the back side for 12 sec, directly plunge-frozen into liquid ethane cooled by liquid nitrogen bath, transferred to liquid nitrogen and clipped into an Autogrid assembly (Thermo Fisher Scientific) for further handling. The grid with the cells was stored in liquid nitrogen until further use. Note that the holey gold film covered TEM grids demonstrated better mechanical stability during the blotting procedure compared to holey carbon film covered grids (QUANTIFOIL®, Quantifoil Micro Tools), resulting in a larger intact film area with retained cells.

The frozen samples were transferred, under cryogenic conditions, using a VCM/VCT500 cryo -transfer system (keeping the sample at <-170°C, Leica Microsystems), into an ACE600 cryo-coater (Leica Microsystems). The samples were sputter-coated with a ∼20 nm platinum conductive protection layer^54,86^, while being kept at <-153°C. The coated samples were transferred to a cryo -fluorescence upright wide-field microscope (THUNDER Imager EM Cryo CLEM, Leica Microsystems) equipped with a 50x objective possessing a high NA of 0.90 (HC PL APO, CRYO CLEM, Leica Microsystems; ∼373 nm XY-resolution and ∼679 nm Z-resolution at 550 nm wavelength) and a cryo-stage (temperature set to -180°C), which accommodates up to two TEM Autogrids. The sample grid plane is oriented perpendicular to the optical axis during imaging within the THUNDER cryo-light microscope (cryo-LM).

Grid overviews (**Supplementary Fig. 2a**) in reflected light (RL; using a LED3 fluorescence light source (Lumencor) and a longpass filter >425 nm) and FL (with 450 -490 nm excitation and 500-550 nm emission filters) channels were obtained within the THUNDER cryo -LM using the embedded Navigator module of LAS X software package (Leica Microsystems) by stitching individual fields of view (271.08 µm x 271.08 µm). Zones with cells exhibiting EYFP FL and an intact support film were investigated in RL, FL and transmitted light (TL; using light of a halogen lamp) regimes and regions of interest (ROIs) containing suitable crystals were identified (see **Supplementary Fig. 2b**).

For each ROI containing a crystal of interest, a 3D volume was imaged as a ∼40 µm dual-channel Z-stack using RL and FL channels (**Supplementary Fig. 3**) and a system-optimized Z-step size of 0.445 µm. THUNDER deconvolution^87^ technique embedded in the LAS X software was applied to the FL Z-stack data to improve the contrast of the FL images. This dual-channel RL/FL Z-stack was used to locate the 3D center of the crystal (X_c_, Y_c_, Z_c_), as shown in **Supplementary Fig. 3**. Using the FL data of the Z-stack, the position (X_c_, Y_c_, Z_c_) was determined as the center of the negative FL signal of the crystal (**Supplementary Fig. 3f**). The Z_c_-depth of the crystal’s center was calculated relative to the Z-position of the top surface of the sample (Z_s_) at (X_c_, Y_c_) location. The Z_s_-position was determined using the RL Z-stack data and set as the Z = 0 reference (**Supplementary Fig. 3e**). The determined lateral position of the crystal center (X_c_, Y_c_) was marked with a dot on the maximum intensity Z-projection of the RL channel of the ROI containing the crystal (see **Supplementary Fig. 3c** and **Supplementary Fig. 4b**). This RL image and the Z-depth of the crystal’s center (Z_c_) were used for subsequent image correlation and crystal localization procedures as depicted in **Supplementary Fig. 4b and** **Supplementary Fig. 5.**

Note that the metallic Pt sputter coating of the samples, in addition to mitigating charging and protecting against unwanted ion beam damage during subsequent FIB/SEM imaging^54,86^, also amplified the light reflection from the coated top surface of the samples. This enhanced the surface imaging contrast for RL microscopy and improved FIB/SEM surface imaging, thereby facilitating both the determination of the Z_s_-position of the sample top surface (**Supplementary Fig. 3c, e, g**) and subsequent correlation between FIB and RL images inside the FIB/SEM microscope (**Supplementary Fig. 4b**). We observed that ∼20 nm thick Pt sputter layers provided good light reflection while remaining sufficiently transparent for TL and FL signals, while thicker layers (*e.g.* ∼40 nm) significantly dampened FL imaging.

Grids containing the crystals of interest were unloaded from the cryo -light microscope (cryo-LM) and, using a VCM/VCT500 cryo-transfer system (<-170°C, Leica Microsystems), placed into a 40° pre-tilted cryo-FIB/SEM grid holder (Leica Microsystems), which accommodates up to two TEM Autogrids per session, and inserted into an Amber cryo-FIB/SEM microscope (Tescan) equipped with a passively cooled cryo -stage (maintained at <-143°C, Leica Microsystems). The angle between the ion and electron beams is 55° for the Amber FIB/SEM microscope. To avoid unwanted sample damage during navigation and monitoring of the milling process, SEM and FIB imaging were performed with 0.5 kV electrons of 10 pA current and with 30 kV Ga^+^ ions of 10 pA current throughout the described cryo -FIB milling procedure.

In the cryo-FIB/SEM microscope, an overview SEM image of a TEM grid was acquired with the electron beam at a 65° angle to the grid plane (the default angle for the Autogrid-compatible sample holder used). To enable rapid localization of ROIs containing crystals of interest, this overview SEM image was correlated with the combined RL and FL maps derived from cryo-LM imaging (**Supplementary Fig. 4a**). This was performed using a three-point image correlation procedure (based on image affine transformations) of the CORAL module within the Essence software (Tescan) which controls the FIB/SEM instrument. Subsequently, for each ROI containing a target crystal, the following steps were performed.

An ROI with a crystal of interest was positioned at the intersection of the SEM and FIB beam foci, and the sample was rotated such that the ion beam was at a 90° angle to the sample grid plane. The ROI was then imaged by FIB, and the resulting image was correlated, using the three-point correlation procedure, with the previously obtained LM image (containing the marked lateral position of the crystal’s center (X _c_, Y_c_); see **Supplementary Fig. 4b**). Using the DrawBeam module of the Essence software, fiducial markers were drawn typically as two 2 µm × 5 µm rectangles separated by a 12 µm gap on the correlated overlay of FIB and LM images, positioning the crystal’s center midway between the fiducials ( **Supplementary Fig. 5a, b**). These fiducial markers were milled (**Supplementary Fig. 5c**) by raster scanning over the drawn pattern using a 30 kV ion beam with a 50 pA current to a depth of ∼1 µm. For the calculations herein, we used an experimentally derived cryo-FIB milling rate of ∼4 µm³/nA/s. Subsequently, the ROI was positioned such that the ion beam was at a 20° angle to the sample plane and then imaged by FIB. The position of the crystal’s center at the 20° angle FIB view was estimated to be Z_c_ × cos 20° lower than the top surface above the crystal center (Z = Z_s_) defined at this FIB view by the previously milled two fiducial markers (see **Supplementary Fig. 5d, e, f**). The third fiducial marker (a 3 µm × 12 µm rectangle) was drawn 3.25 µm lower than the estimated position of the crystal’s center and FIB-milled at 150 pA current to a depth of ∼2 µm. All three fiducial markers per each crystal served as 3D locators of estimated crystal center positions for subsequent procedures.

After creating all the fiducial markers for all the crystals of interest, the samples were coated with a protective (see *e.g.*^41,45,54^) organoplatinum layer using a gas injection system (Pt-GIS, 60°C, 45 s, ∼400 nm/min layer deposition rate) integrated into the Amber cryo-FIB/SEM microscope. The resulting protective Pt-GIS layer was ∼300 nm thick. For each crystal of interest, the following FIB milling steps were performed.

A center of a crystal of interest was relocalized at different FIB view angles using the previously FIB-marked fiducials. "Rough" cryo-FIB milling was performed via raster scanning with a 30 kV Ga^+^ ion beam, first, at 90° and then at 20° to the TEM grid plane using high currents of 2.5 nA and 1 nA, respectively, to remove the bulk of the sample as depicted in **Supplementary Fig. 6**. This produced a ∼4.5 µm thick lamella at a 20° angle to the grid plane. Subsequently, a "gentle" cryo-FIB milling procedure was performed by raster scanning with the 30 kV Ga^+^ FIB at 20° to the TEM grid plane in a stepwise manner, reducing the beam current as the lamella was thinned to its final thickness, as shown in **Supplementary Fig. 7**. Milling currents of 150 pA, 50 pA, and 10 pA were used (in this order) to generate the final lamella by decreasing the thickness of the sample to ∼1.5 µm, ∼0.8 µm, and the final thickness, respectively. Such low currents were used for the "gentle" procedure to retain the maximum possible degree of crystalline order of the *in cellulo* crystal (see *e.g.*^45,86^). Lamellae produced for 200 kV and 300 kV ED experiments were ∼4 µm wide and with final thicknesses, corresponding to the reported optimum^40,41^, of ∼250 nm and ∼300 nm, respectively, and were angled at 20° with respect to the TEM grid plane.

Using the VCM/VCT500 cryo-transfer system, the grids were then transferred from the cryo -FIB/SEM microscope to cryo-EM grid boxes (Thermo Fisher Scientific) for storage in liquid nitrogen until further use in ED experiments.

#### Electron diffraction data collection, data processing and structure determination

For 200 kV ED experiments, we used a JEM-F200 cryo-TEM (JEOL) equipped with a cold cathode field-emission electron gun (CFEG), a side-entry dual Autogrid cryo-holder (-177°C, Model 210, Simple Origin), and a scintillator-based CMOS detector (4096 × 4096, 15.5 × 15.5 µm^2^ pixel, TemCam-XF416, TVIPS). Crystal lamellae were identified in imaging mode (**Fig. 3d**), and ED experiments were performed with parallel electron beam illumination at a flux of ∼0.04 e/Å²/s. ED data were collected either from the parallel beam illumination area, which was ∼4.8 µm in diameter on the sample plane, or from a crystal lamella zone of ∼1 µm in diameter, as defined by a selected area aperture. ED data were recorded as still images with the XF416 camera, using 2 × 2 binning and a 2 s exposure. The effective sample-to-detector distance was ∼2310 mm, as calibrated with a standard evaporated aluminum grid (Ted Pella). Adxv ^88^ and ImageJ^85^ software were used for the visualization of the ED images.

For 300 kV ED experiments, we used a Titan Krios G1 cryo-TEM (Thermo Fisher Scientific), equipped with a Schottky field emission gun (XFEG), a cryogenic sample Autoloader (∼-193°C), a GIF BioQuantum energy filter (Gatan/Ametek) and a K3 direct electron detector (5760 × 4092, 5 × 5 µm^2^ pixel, Gatan/Ametek). The sample grids were cryo-transferred from storage to the Autoloader cassette, ensuring the microscope’s rotation (α-tilt) axis was orthogonal to the FIB milling direction. In the cryo -TEM, the crystal lamellae were identified in imaging mode (**Supplementary Fig. 8**). ED data acquisition employed a parallel electron beam with a 14 µm diameter illumination area on the specimen plane and a low electron flux of ∼0.0009 e/Å²/s. The energy filter slit was set to select only electrons with energy loss less than 10 eV. Following the manufacturer’s recommendations (Gatan/Ametek) for 3D ED data collection, the K3 detector was operated in standard electron counting mode (without correlated double sampling; internal frame rate of 1.5 kHz) at a dose rate below 40 e/pixel/s for ED peaks (to avoid significant coincidence loss ^89^–^91^). This dose rate was achieved by setting up the low flux of the parallel electron beam (∼0.0009 e/Å²/s) and validated by acquiring 1 s exposure ED still images from a crystal lamella prior to 3D ED collection. The effective sample-to-detector distance was ∼905 mm, as calibrated with a standard evaporated aluminum grid (Ted Pella). 3D ED data were recorded in a continuous rotation mode collecting ED signal from crystal lamella zones with diameters of 2.5 µm (referred to as “microvolume”) or 1.75 µm (referred to as “nanovolume”) as defined by selected area apertures of the cryo-TEM. The K3 camera recorded at 1 fps during the data collection. Microvolume data were collected during 325 s exposure at a rotation speed of 0.249°/s, covering a total angular wedge of 80.9° (from −40.4° to +40.5° α-tilt) at a fluence of ∼0.3 e/Å². Nanovolume data were collected during 660 s exposure at a rotation speed of 0.1224°/s, covering a total angular wedge of 80.8° (from −40.4° to +40.4° α-tilt) at a fluence of ∼0.6 e/Å². Absorbed radiation doses were estimated using RADDOSE-ED software^57^. The electron-counting data images were saved as 8-bit unsigned TIFF (Tagged Image File Format) stack without detector gain normalization being applied. For further processing, the images were gain-normalized and converted to the CBF (Crystallographic Binary File) format using in-house developed Python 3.12.4 script and 2cbf program of XDS software package^92^.

The 3D ED data were indexed and integrated using XDS software ^92^, and then scaled and merged using AIMLESS software^93^. Initial phases were obtained by molecular replacement with Phaser^94^ using the *Nc*HEX-1 structure, previously obtained using crystals grown by a sitting drop vapor diffusion method, as a search model (PDB 1KHI). *Mg*HEX-1 crystal structure refinements for all collected datasets were carried out using REFMAC5^95^ (version 5.5.0051; electron scattering factors were set with the keyword “source EC MB”) and iterative cycles of manual model building in Coot^96^ (version 0.9.7). PyMOL Molecular Graphics System (version 4.5.0, Schrödinger, Inc.) was used for graphical illustrations.

#### Serial X-ray diffraction data collection, data processing and structure determination

Sample preparation for X-ray diffraction measurements was performed as previously described ^82^. In brief, High Five cells carrying *Mg*HEX-1 protein crystals were harvested into a 1.5 mL tube and allowed to settle by gravitation. In a 90% humidity air stream 0.5 μL cells from the loose pellet were loaded onto a MicroMesh (700/25; MiTeGen) previously mounted on a B1A goniometer base (MiTeGen), which was positioned by a custom-made holder in the optical focus of a standard upright cell culture light microscope. The excess medium was removed using an extra fine liquid wick (MiTeGen). For cryo -protection, 0.35 µL of a 40% PEG200 solution diluted in ESF921 cell culture medium were pipetted onto the cells, again followed by removal of the excess liquid. Cell-loaded MicroMeshes were flash frozen and stored in liquid nitrogen until further use in X-ray diffraction experiments.

X-ray diffraction data collection was performed at the EMBL microfocus beamline P14 at the PETRAIII storage ring (DESY, Hamburg) using the mxCuBE v2 user interface ^97^. The cell-loaded MicroMeshs were mounted on a mini-kappa goniostat attached to an MD3 diffractometer using the MARVIN Sample Changer. Using a 7 x 3 µm² double focussed X-ray beam characterized by an energy of 12.7 keV and a flux of 1.71 x 10^13^ ph/s MicroMeshes were scanned employing a previously established serial helical grid scan strategy^2,23^ to collect complete datasets using the EIGER 16M detector (DECTRIS) at 100 K. During 20 ms irradiation at each position the MicroMesh was rotated by 1°. Absorbed radiation doses were estimated using RADDOSE-3D software^57^. Data collection parameters are presented in **Table 1**.

To process X-ray diffraction data using CrystFEL^98^, a geometry file specifying information about the sample to detector distance (clen), the wavelength (photon_energy), the size of the detector (max_fs and max_ss) as well as the beam position on the detector relative to the detector corner (corner_x and corner_y) is required. Then peak detection parameters can be specified using the graphical user interface of CrystFEL version 10.0/10.1 or the check-peak-detection script of version 9.1 and older versions using peak finding algorithm peakfinder8. In general, for this study, a threshold of 0, a local_bg_radius of 3, min_res of 50, a max_res of 2200 and a max_pix value of 50 were a suitable starting point. The min_snr and min -pix were adjusted to allow proper peak detection for each sample. Afterwards, all frames were indexed by Moslfm, and the correct lattice and unit cell parameters were optimized, and cycles of beam position refinement were performed by executing the detector_shift script (version 9.1) on the obtained stream files. If the optimized beam position did not differ more than 0.1 mm from the input, the beam position was accepted and all frames were indexed invoking mosflm-latt-cell, mosflm-latt-cell, xds and xgandalf. Then the peakogram-stream script was executed to set the maximum allowed intensity for each resolution range to separate salt reflections from protein reflections applying a filtering script. Finally, the filtered intensities were scaled and merged. The resolution limit was set where CC* is above 50 %, SNR above 0.5 and the completeness over 95%. Mtz-files for subsequent modeling were generated using get_hkl. To set up peak detection parameters CrystFEL version 10.0 was applied, for indexing CrystFEL version 10.1 or 10.2, while for merging and scaling CrystFEL version 9.1 or 10.2 were used.

Phases were retrieved by molecular replacement with Phaser^94^ using the *Nc*HEX-1 structure as a search model (PDB 1KHI). Structure refinements for all generated datasets were carried out using PHENIX ^99^ (version 1.19.2-4158) and iterative cycles of manual model building in Coot^96^ (version 0.9.7). Simulated-annealing omit maps were calculated in PHENIX. Applying the FastFourierTransform program, electron density maps with CCP4 extensions were saved and loaded in the PyMOL Molecular Graphics System (version 4.5.0, Schrödinger, Inc.) for graphical illustrations of contoured omit maps.

### Data availability

Protein Data Bank: The atomic coordinates and structure factors of the SSX structure as well as of the microvolume and nanovolume 3D ED/MicroED structures of *Mg*HEX-1 have been deposited in the RCSB Protein Data Bank (PDB) with accession code 8QLX [https://doi.org/10.2210/pdb8QLX/pdb], 9RVB [https://doi.org/10.2210/pdb9RVB/pdb] and 9RVD [https://doi.org/10.2210/pdb9RVD/pdb], respectively.

### Author contributions

V.P. and L.R. conceived the *IncellED* approach, which was designed with Š.B., D.P., K.K., J.H., T.B., Z.F., and R.T.. Š.B. performed *in cellulo* crystallization experiments, supervised by R.T., Z.F., and L.R.. Light microscopy, sample blotting, FIB/SEM, and TEM experiments were performed by Š.B., D.P., K.K., V.P. with the help of T.B., R.T., Z.F., and were generally overseen by V.P.. K.K. and V.P. processed the ED data and performed molecular replacement with the help of J.H.. L.R. and V.P. refined the ED structures, and calculated the electrostatic potential maps. Supervised by L.R., J.B. performed SSX experiments, processed the SSX data, performed molecular replacement, refined the X-ray structure, and calculated the electron density map. The manuscript was prepared by Š.B., D.P., K.K, J.H., R.T., L.R. and V.P. with input, discussions and improvements from all authors.

## Acknowledgements

The authors would like to thank Sahil Gulati (Gatan/Ametek) and Samuel Záchej (Tescan) for helpful technical discussions regarding the K3 camera operation and the FIB/SEM microscope operation, respectively. We acknowledge the Electron Microscopy Core Facility, IMG, Prague, Czech Republic, supported by the MEYS CR (LM2023050) and the European Regional Development Fund (No. CZ.02.1.01/0.0/0.0/18_046/0016045, No. CZ.02.01.01/00/23_015/0008205), for their support with experiments and data collection performed at cryo-LM, FIB/SEM and 200 kV TEM microscopes. We acknowledge Cryo-electron microscopy and tomography core facility CEITEC MU of CIISB, Instruct-CZ Centre, supported by the MEYS CR (LM2023042) and the European Regional Development Fund (Project “Innovation of Czech Infrastructure for Integrative Structural Biology”; No. CZ.02.01.01/00/23_015/0008175), for their support with 300 kV TEM experiments. We thank Jiří Nováček for his assistance with these experiments. The synchrotron diffraction data was collected at the P14 beam line operated by EMBL Hamburg at the PETRA III storage ring (DESY, Hamburg, Germany). We thank Gleb Bourenkov for the assistance in using the beamline. Š.B., T.B., Z.F. and R.T. have been supported by the European Regional Development Fund (No. CZ.02.1.01/0.0/0.0/15_003/0000441). J.H., K.K. and V.P. acknowledge the support from the European Regional Development Fund (“Structural dynamics of biomolecular systems (ELIBIO)"; No. CZ.02.1.01/0.0/0.0/15_003/0000447). K.K. and V.P. also thank the HORIZON EUROPE framework program for research and innovation (Grant Agreement No. 101094299 “IMPRESS”). L.R. thanks the German Federal Ministry for Education and Research (BMBF; grant 05K18FLA) for the support.

## Supplementary Figures

**Supplementary Figure 1.**
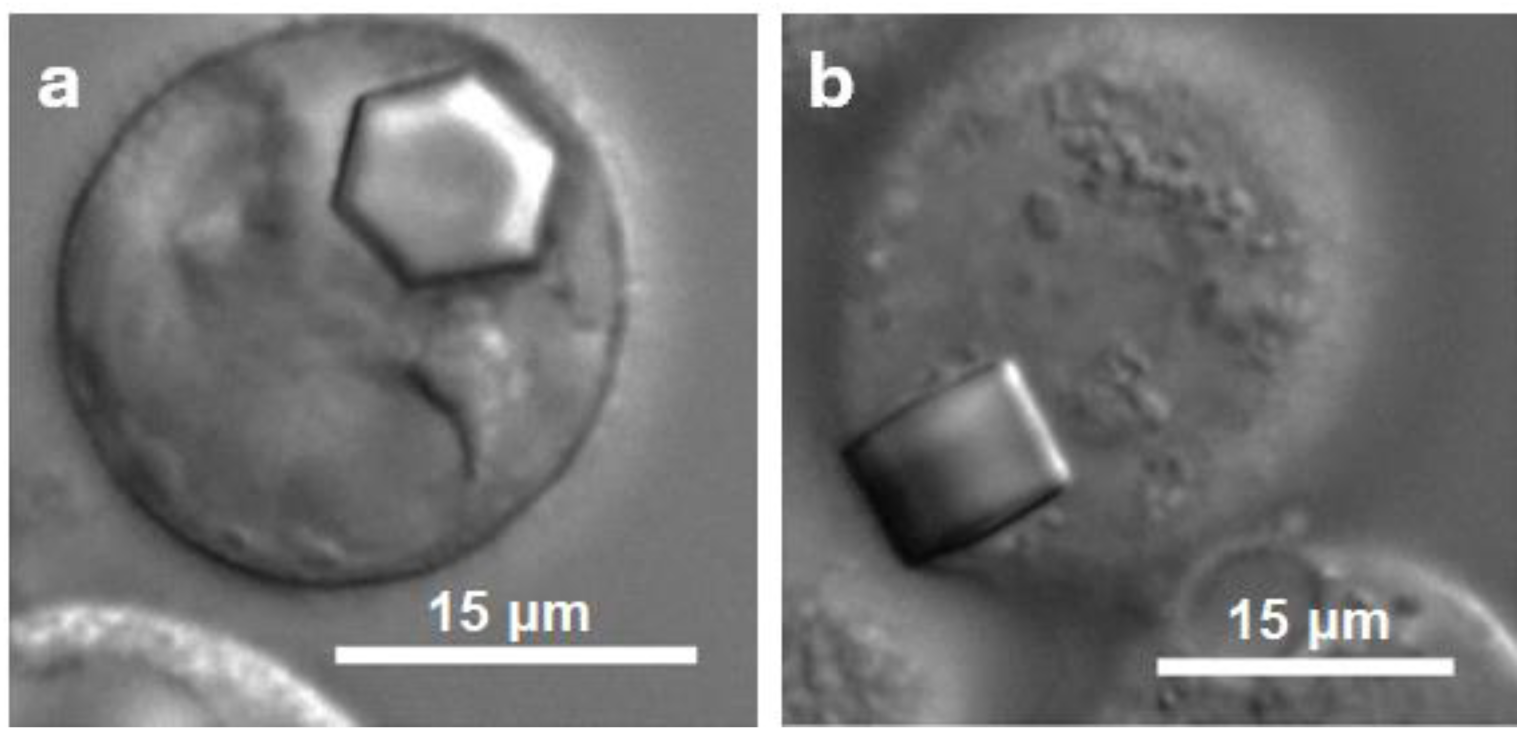
Light microscopy images of ordered structures detected in High Five insect cells four days after infection with an rBV encoding *Mg*HEX-1. The predominant morphology is a broad rectangle with a hexagonal cross-section. Cells were imaged using differential interference contrast (DIC)^1^ and a 100x objective on a Nikon Ts2R-FL. (**a**) A front view showing the hexagonal cross-section of an *in cellulo Mg*HEX-1 crystal. (**b**) A rectangular side view of an *in cellulo Mg*HEX-1 crystal.

**Supplementary Figure 2.**
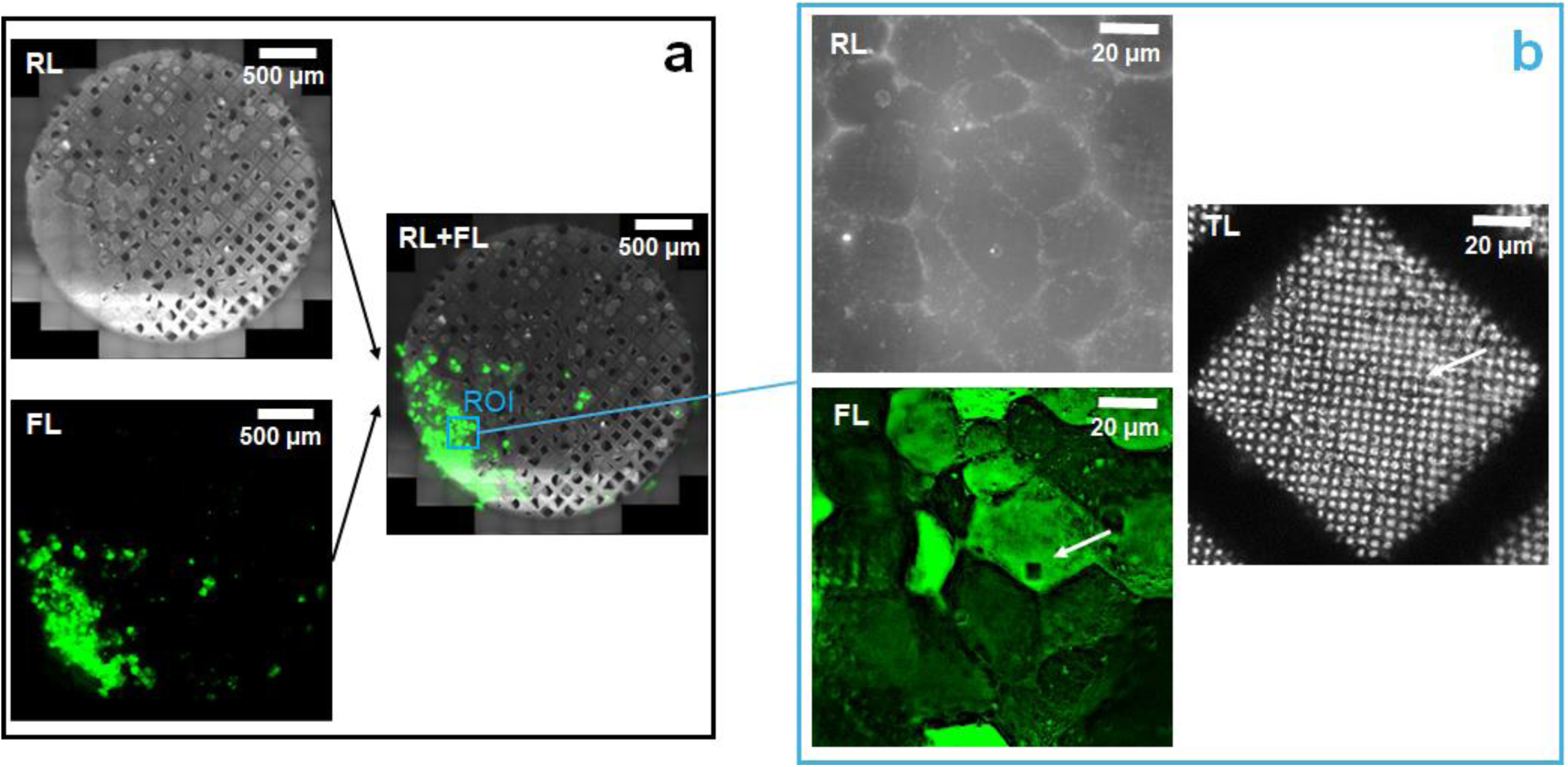
Cryo-LM investigation of a TEM grid with plunge-frozen High Five insect cells containing *Mg*HEX-1 crystals. (**a**) An overview of the grid acquired using reflected light (RL) and fluorescence (FL) channels, with the overlay (RL+FL). (**b**) A region of interest (ROI) containing an *in cellulo Mg*HEX-1 crystal using RL, FL, and transmitted light (TL) imaging. A crystal of interest (indicated by a white arrow) was identified via FL and TL images. The crystals selected for subsequent *IncellED* processing were located on an intact support foil (as confirmed by RL and TL imaging). *NOTE: due to the weak ability of Ga^+^ cryo-FIB to mill through the metal bars of TEM grids, the chosen crystals were also located at least 15 µm (as deduced from TL images) from the ∼22.5 µm thick grid bars. This ensured that, during subsequent 3D ED single-crystal data collection within a cryo-TEM, the electron beam probing a crystal was not blocked by the grid bar over a wide angular tilt, thus providing data completeness sufficient for reliable structure elucidation* ^2–5^. *However, in our experience, plasma Xe^+^ FIBs are much more efficient at milling through the metal grid bars under cryogenic conditions, although this has not yet been tested for the IncellED workflow. Therefore, using these more advanced but less common and available plasma FIBs*^6–8^ *could increase the number of crystals on the TEM grid suitable for broad angle wedge 3D ED/MicroED data collection. Alternatively, merging 3D ED datasets with smaller angle wedges and lower completeness, that are collected from multiple crystals near the TEM grid bars, could also be a solution, similar to what is mentioned in Discussion when addressing the low-symmetry/completeness issue*.

**Supplementary Figure 3.**
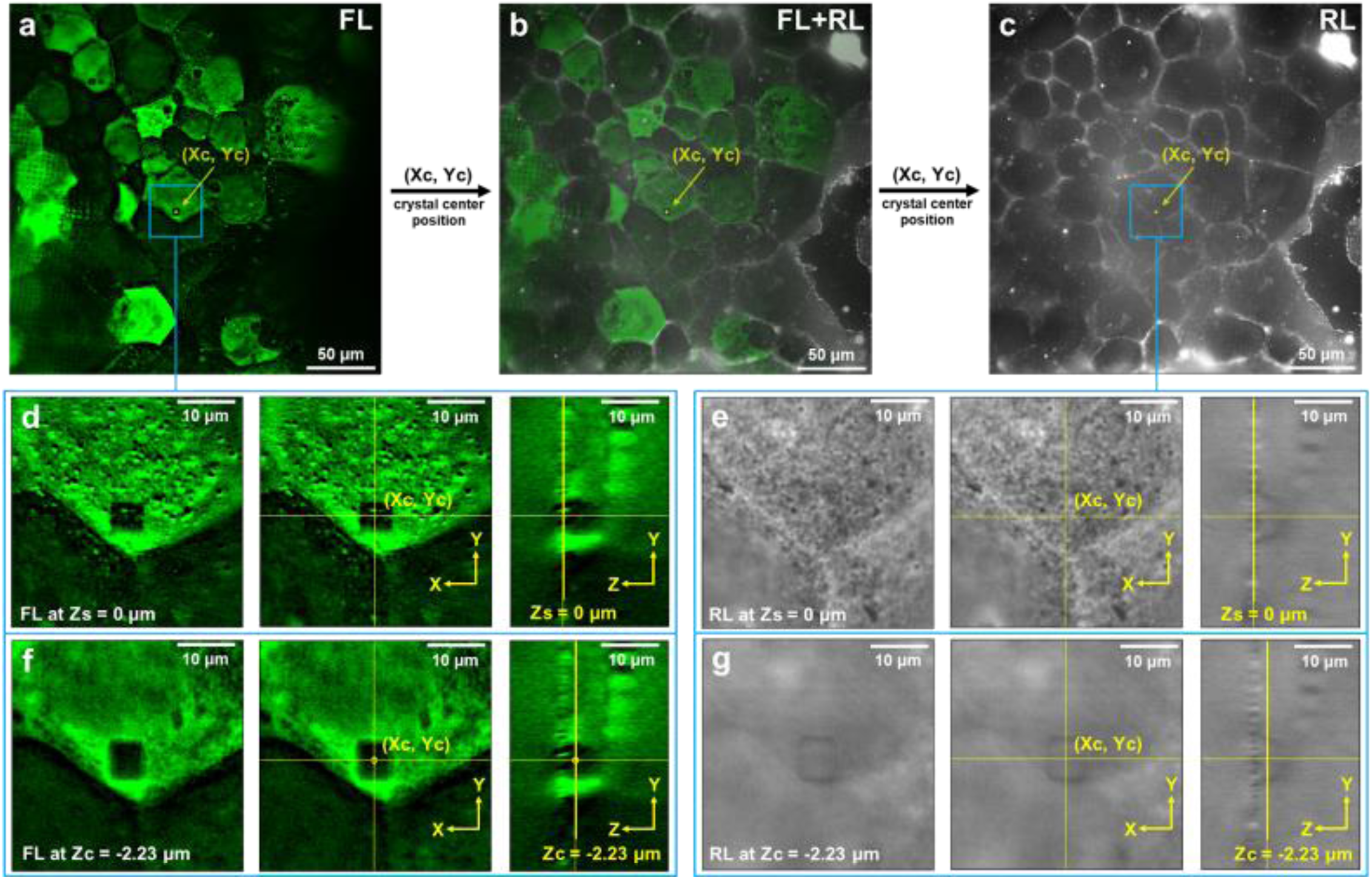
3D dual-channel RL/FL cryo-LM imaging of an ROI, containing an *in cellulo* crystal of interest, to determine its 3D center position (X_c_, Y_c_, Z_c_). A Z-stack of the ROI from **Supplementary Fig. 2b** was acquired using both RL and FL modes, inherently correlated in XYZ by the microscope system, with a Z-step of ∼0.445 µm. (**a**) shows the FL Z-slice of the ROI around the crystal’s center (Z_c_). (**b**) presents an overlay of (**a**) with the maximum intensity Z-projection of the RL imaging Z-stack, which is displayed in (**c**). The lateral position of the crystal’s center (X_c_, Y_c_) is indicated by a yellow dot in (**a**), (**b**), and (**c**). For a smaller region indicated by a blue square in (**a**) and (**c**), orthogonal views of the FL and RL Z-stacks are shown for different Z-slices: at the top sample surface (Z_s_) and (X_c_, Y_c_) position - FL in (**d**), RL in (**e**); and at the crystal’s center (Z_c_) - FL in (**f**), RL in (**g**). The 3D position of the crystal’s center (X_c_, Y_c_, Z_c_) was determined as the center of the area lacking the EYFP-derived FL signal as seen in (**d**) and (**f**). The reference Z-level (Z=0) was assigned to the top sample surface (Z_s_) at (X_c_, Y_c_) and identified using the RL Z-stack as depicted in (**e**) and (**g**). The lateral position (X_c_, Y_c_) of the crystal’s center, initially determined from the FL data, was transferred to the RL Z-stack via the overlay in (**b**). ImageJ software^9^ was used to perform dual-channel RL/FL Z-stack manipulation, maximum intensity Z-projection, and Z-stack orthogonal view representation.

**Supplementary Figure 4.**
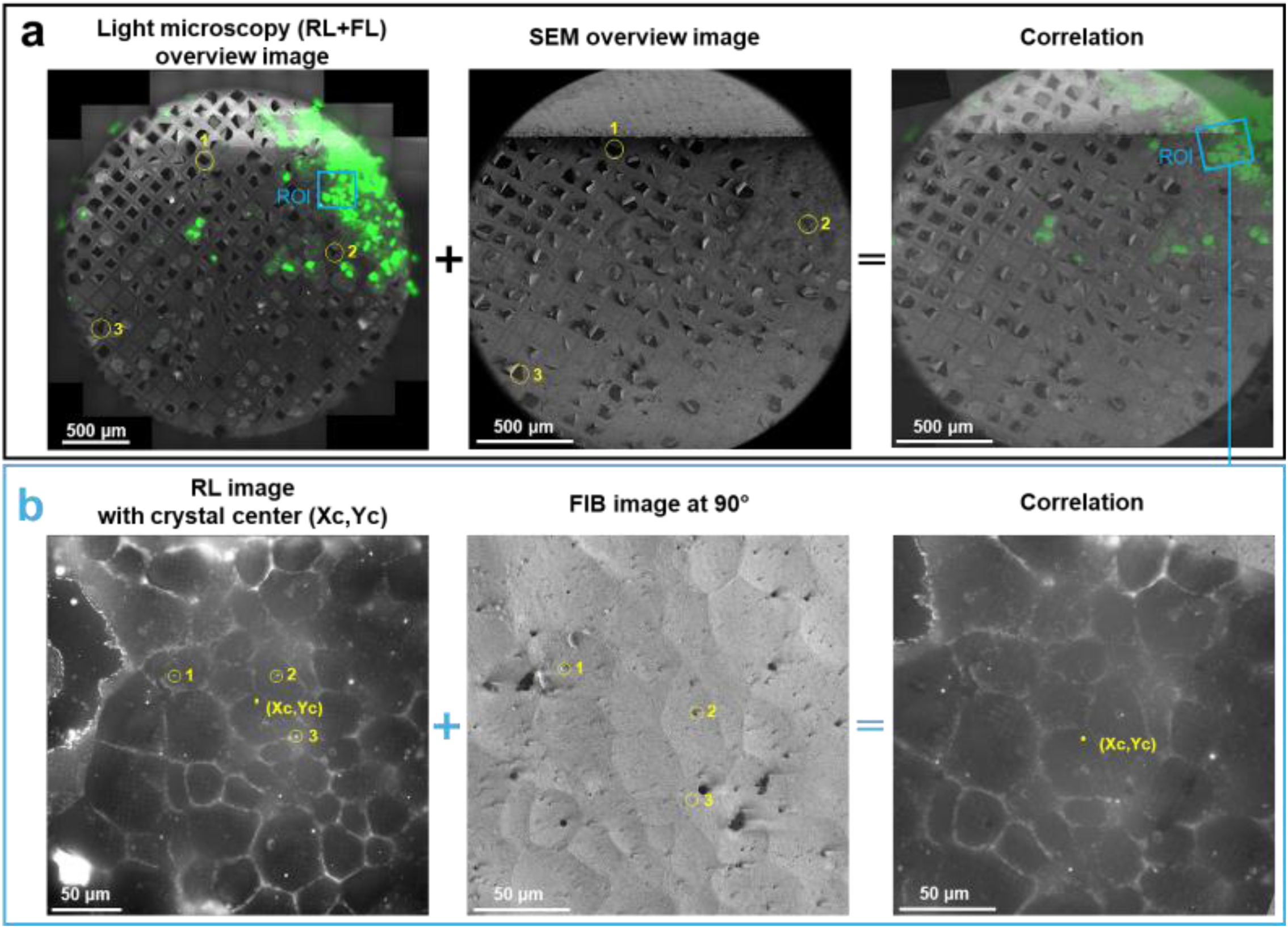
Lateral (XY) localization of an *in cellulo* crystal of interest on a TEM grid within a cryo-FIB/SEM microscope. Panel (**a**) shows the preliminary localization of the ROI containing the crystal using an SEM overview image of the TEM grid and the overlaid RL/FL overview obtained with cryo -LM (identical to **Supplementary Fig. 2a** but rotated by 180° for convenience). The RL/FL image with the ROI labelled by a blue square was aligned with the SEM image using a three-point correlation procedure embedded in the cryo-FIB/SEM microscope’s software. Three common features - foil cracks and metal grid bars, indicated by numbered yellow circles in (**a**) - were seen in both the SEM image and the LM image and used as reference points for the correlation. This correlation guided the finding of the ROI within the FIB/SEM microscope, and the region containing the crystal of interest was then imaged with a 30 kV Ga^+^ FIB at 90° to the TEM grid plane. Panel (**b**) presents the localization of the crystal of interest using this FIB image and the maximum intensity RL Z-projection image previously acquired with cryo-LM (identical to **Supplementary Fig. 3c** but rotated by 180° for convenience). The RL image with the marked position ((X_c_, Y_c_), a yellow dot) of the crystal center was aligned with the FIB image using the three-point correlation procedure and common features (indicated by numbered yellow circles in (**b**)) of the sample surface identified by both the RL and the 90° FIB cryo-imaging. Typically, such reference points include surface contamination (*e.g.*, ice nano-/micro-crystals), surface irregularities, cracks, and patterns of the TEM grid support film. Especially when enhanced by a metallic Pt sputter coating, these features serve as effective fiducials because they both scatter light well and are clearly identifiable in FIB imaging. *NOTE: these "natural" fiducials, inherent to the TEM grid preparation process, require visual identification and manual selection, which can be a time-consuming process. We believe that the recent interest (see e.g.*^10^*) in using AI-driven approaches for correlative light-electron microscopy (CLEM) could facilitate and accelerate this step of the IncellED workflow. Future improvements could also involve artificially fabricated fiducial markers. For example, this might include utilizing specifically patterned TEM grids for sample cryo-preparation, such as finder EM grids*^11^*, or applying special FIB/embedding patterning to cryo-samples (see e.g.*^12,13^*) prior to acquiring cryo-LM data*.

**Supplementary Figure 5.**
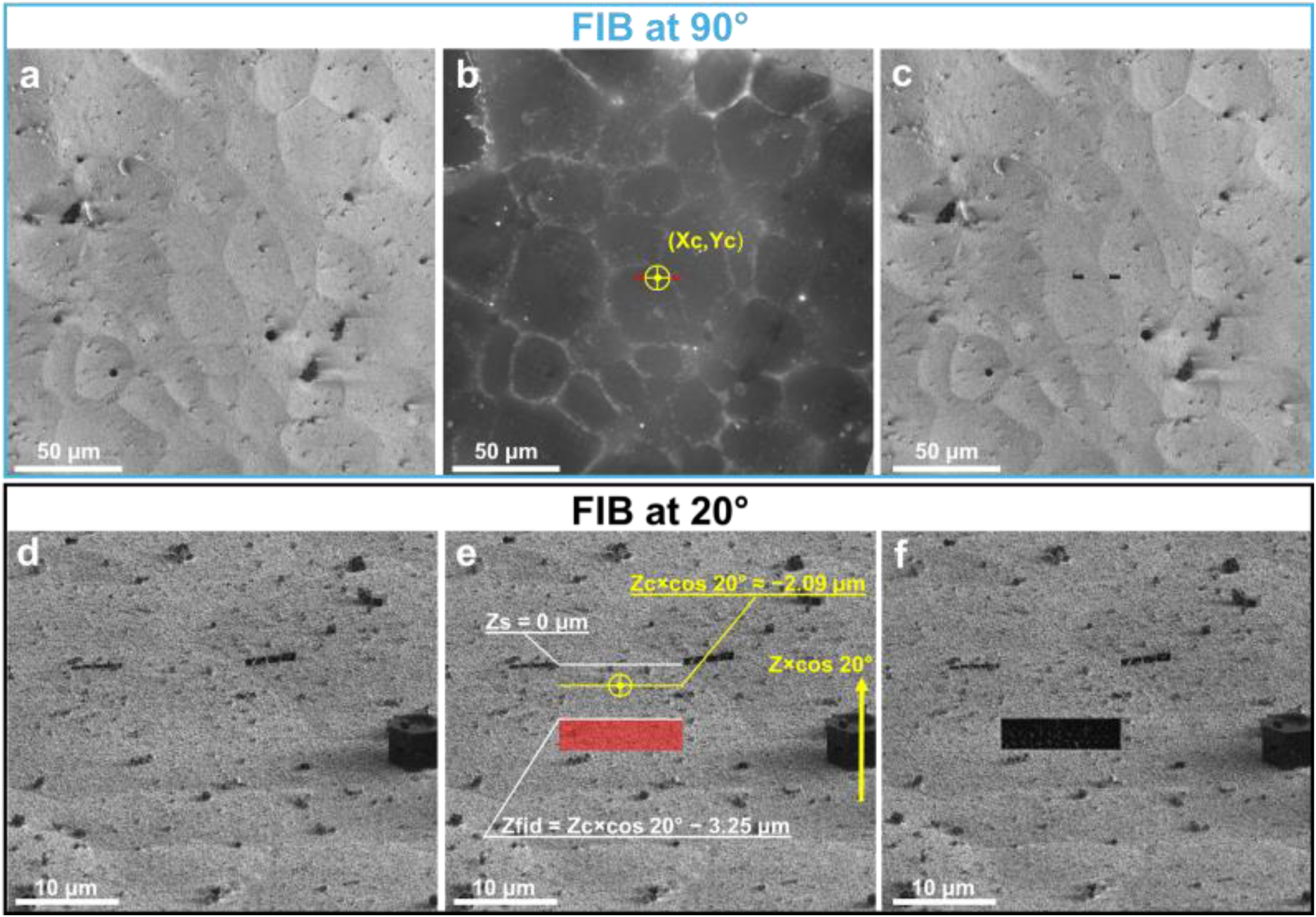
Creation of fiducial markers by a 30 kV Ga^+^ FIB to indicate the center of a target crystal and to guide subsequent lamella production by cryo-FIB milling. FIB at a 90° angle to the sample grid plane: (**a**) FIB image of the ROI containing the crystal of interest before fiducial marker creation; (**b**) a pattern for the fiducial markers was drawn as two 2 µm × 5 µm red filled rectangles separated by a 12 µm gap, with the crystal’s center (indicated by the center of the yellow crossed circle, 12 µm in diameter) positioned midway between these markers; this drawing was performed on the correlated overlay of this FIB image and the LM image, which contained the lateral position (X_c_, Y_c_) of the crystal’s center (note that (**a**) and (**b**) are identical to the images from **Supplementary Fig. 4b**); (**c**) FIB image of the ROI containing the target crystal after fabricating these two fiducial markers FIB rastering over the drawn patterns. FIB at a 20° angle to the sample grid plane: (**d**) FIB image of the ROI containing the crystal of interest before fabricating a third fiducial; (**e**) the sample top surface (X_c_, Y_c_, Z_s_ = 0) at the position of the crystal center was localized midway between two previously created fiducials; the depth of the crystal center Z_c_ ≈ -2.23 µm (**Supplementary Fig. 3d, f**) is projected as Z_c_ × cos 20° ≈ -2.09 µm at the 20° FIB view; a pattern for the third fiducial marker was drawn as a 3 µm × 12 µm red filled rectangle placed 3.25 µm lower (Z_fid_) than the estimated position of the crystal’s center (indicated by the center of the yellow crossed circle); (**f**) FIB image of the ROI containing the target crystal after fabricating the third fiducial marker by FIB rastering over the drawn pattern, which indicated the Z-position of the crystal’s center. *NOTE: the approximation of the sample top surface in (**e**) as a straight line (Z_s_=0; white line) was valid, as the surface was sufficiently flat (with regard to the sub-1 µm Z-resolution of the microscope’s objective) within the 12 µm range between the fiducials (see e.g.* ***Supplementary*** Fig. 3e, g*). This was common for crystals investigated in the study. Otherwise, the shape of the surface around the crystal of interest, obtained via RL imaging, should be considered, which might necessitate a fiducial pattern different from the one described here*.

**Supplementary Figure 6.**
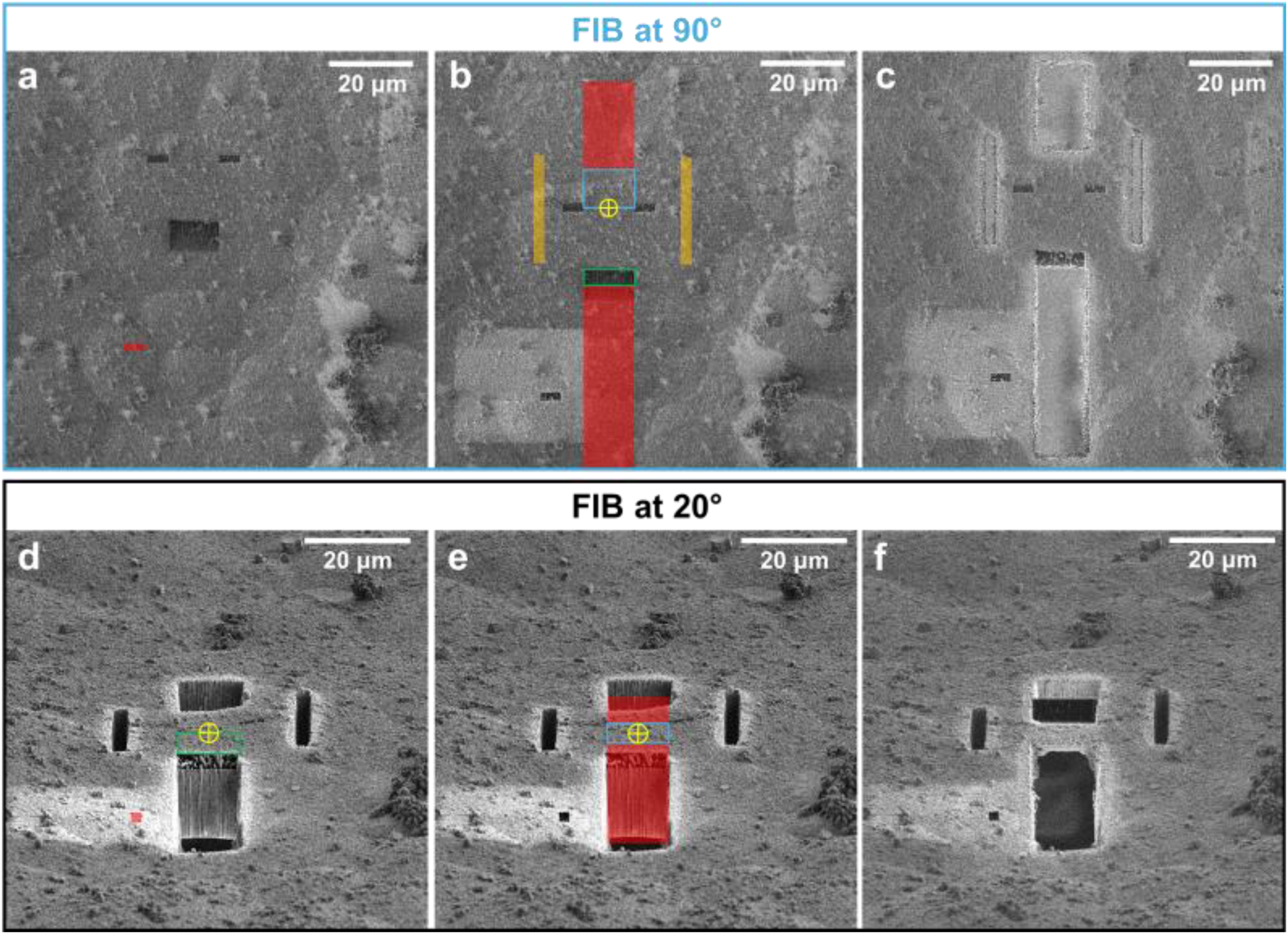
“Rough” cryo-FIB milling step for the preparation of a cellular lamella containing a portion of a target crystal. First, the “rough” milling was performed using a 30 kV Ga^+^ FIB at a 90° angle to the sample grid plane: (**a**) FIB image of the ROI before milling; a red filled rectangular pattern on the side of the target was drawn, pre-milled with the 50 pA ion beam, and used as a fiducial to assist drift correction during the subsequent “rough” milling at the 90° angle; (**b**) using the previously created fiducials (**Supplementary Fig. 5b, c**), the center of the target crystal was re-localized at the 90° FIB view (indicated by the center of the yellow crossed circle, placed midway between two fiducial markers); to prevent ion beam damage to the crystal during the subsequent milling and to preserve the previously made fiducial (which indicates the crystal’s Z-position), a pattern for bulk sample removal was drawn as two 12 -µm-wide red-filled rectangles and placed at least 9 µm from the crystal’s center (see the 9 µm × 12 µm blue rectangle) and away from the fiducial (see the 3.5 µm × 12 µm green rectangle); a pattern for lamella stress-relief gaps (see *e.g.*^14^) was drawn as two 2.5 µm wide orange filled rectangles and placed at least 15 µm away from the crystal’s center; (**c**) FIB image of the ROI after cryo-FIB milling with a high current of 2.5 nA by rastering over the filled rectangular patterns in (**b**). Afterwards, the “rough” milling was continued using the 30 kV Ga^+^ FIB at a 20° angle to the sample grid plane: (**d**) FIB image of the ROI before milling at the 20° angle; using the previously created fiducial marker (**Supplementary Fig. 5e, f**) the crystal’s center (indicated by the center of the yellow crossed circle) was re-localized at the 20° FIB view to be 3.25 µm above the edge of the fiducial (see the 3.25 µm × 12 µm green rectangle); a red filled square pattern on the side of the target was drawn, pre-milled with the 50 pA ion beam, and used as a fiducial to assist drift correction during the subsequent “rough” milling at the 20° angle; (**e**) a pattern for removing the bulk of the sample was drawn as two 12 µm wide red filled rectangles separated by a 4.5 µm × 12 µm blue rectangle with its center placed at the center of the crystal; (**f**) FIB image of the ROI after cryo-FIB milling with a high current of 1 nA by rastering over the filled rectangular patterns in (**b**); the “rough” cryo-FIB milling step resulted in a ∼4.5 µm thick and 12 µm wide lamella at a 20° angle to the grid plane.

**Supplementary Figure 7.**
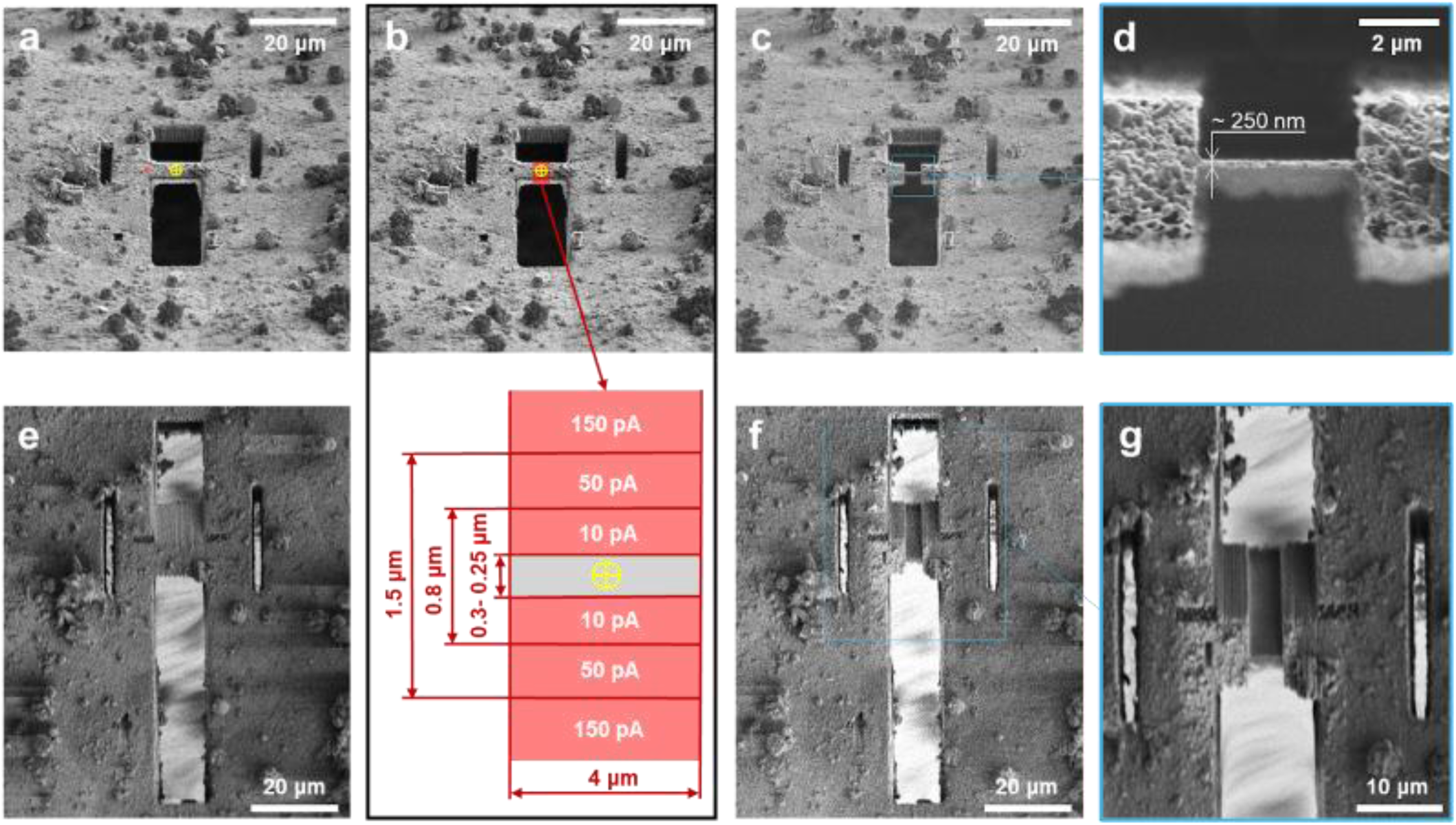
Final “gentle” cryo-FIB milling step for the preparation of a cellular lamella containing a portion of a target crystal. The lamella, typically resulting from the “rough” cryo -FIB milling stage of sample preparation (**Supplementary Fig. 6**), was ∼4.5 µm thick and 12 µm wide, forming a 20° angle to the grid plane. Prior to the “gentle” cryo-FIB milling, micrographs of the lamella were acquired by FIB imaging at a 20° angle to the grid plane (**a**) and by SEM imaging (**e**) (the angle between the ion and electron beams is 55° for the particular FIB/SEM microscope used here). In the FIB image (**a**), the crystal’s center (indicated by the center of the yellow crossed circle) was positioned at the center of the ∼4.5 µm × 12 µm lamella projection at the 20° view. A red filled square pattern was drawn on the side of the target in ( **a**), pre-milled with a 10 pA ion beam, and used as a fiducial to assist drift correction during the subsequent milling. (**b**) shows 4 µm wide red filled rectangular patterns drawn on the FIB image and designed for the “gentle” FIB-milling. This was performed by ion beam rastering over these patterns using low currents of 150 pA, 50 pA, and 10 pA to progressively decrease the sample’s thickness to ∼1.5 µm, ∼800 nm, and a final thickness of ∼300 - 250 nm, respectively. FIB and SEM images of the ∼4 µm wide lamella resulting from the final "gentle" cryo-FIB milling step are presented in (**c**) and (**f**), and with magnified views in (**d**) and (**g**), respectively. This particular lamella of ∼250 nm in thickness was subsequently used for 200 kV ED measurements (see **Fig. 3**). *NOTE: a comparison of* ***Supplementary* Fig. 7a** *to* ***Supplementary* Fig. 6f** *reveals the appearance of ice nano/micro-crystal contamination. In general, after the initial "rough" milling, a sample was either – as in this particular case illustrated here – dismounted, while remaining in the holder, and stored in liquid nitrogen until further use, or milling immediately continued with the "gentle" procedure using low ion currents. Storing the sample may introduce negligible ice crystal contamination in the ROI, which can be quickly removed during the subsequent "gentle" milling. This pause between “rough” and “gentle” milling steps offers a practical advantage for FIB/SEM instrument scheduling in multi-user facilities*.

**Supplementary Figure 8.**
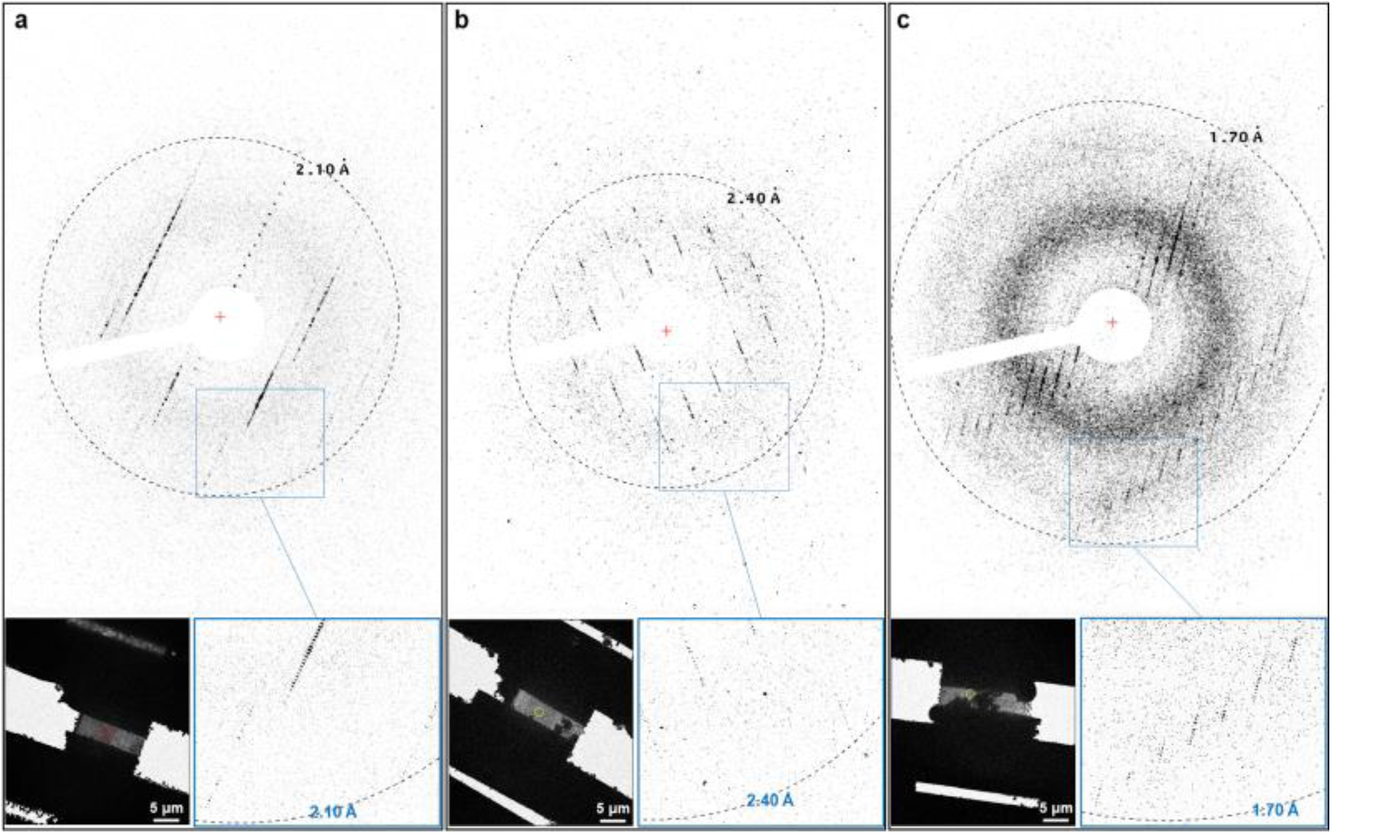
300 kV ED diffraction patterns collected from different ∼300 -nm thick cellular lamellae containing parts of *in cellulo Mg*HEX-1 crystals. The ED signals were registered using the electron-counting DED K3 camera with the electron energy filter slit set to a 10 eV energy loss and the same electron beam fluence of ∼0.0009 e/Å²: (**a**) ED pattern, corresponding to a ∼0.249° wedge, from the continuous-rotation microvolume 3D ED dataset (Table 1); the left inset shows the TEM image of the lamella and the ED collection zone of 2.5 µm in diameter (crystal volume∼1.6 µm³) defined by a selected area aperture of the cryo-TEM and depicted by a red circle; the right inset with the zoomed ED pattern shows the diffraction signal up to ∼2.10 Å; (**b**) ED pattern, corresponding to a ∼0.1224° wedge, from the continuous-rotation nanovolume 3D ED dataset (Table 1); the left inset shows the TEM image of the lamella and the ED collection zone of 1.75 µm in diameter (crystal volume ∼0.8 µm³) defined by a selected area aperture of the cryo-TEM and depicted by a yellow circle; the right inset with the zoomed ED pattern shows the diffraction signal up to ∼2.40 Å; (**c**) ED pattern collected as a still image from a 1.75 µm in diameter area of the crystal lamella (crystal volume ∼0.2 µm³) defined by a selected area aperture of the cryo-TEM and depicted by a yellow circle in the left inset; the left inset shows the TEM image of the lamella; the right inset with the zoomed ED pattern shows the diffraction signal up to ∼1.70 Å. The lamella shown in (**c**) was contaminated by ice microcrystals during cryo-transfer to the cryo-TEM from storage, which prevented the collection of 3D ED data with an angular wedge reasonable for structure determination. Red crosses in (**a**), (**b**), (**c**) ED patterns indicate the position of the electron beam center. Adxv software^15^ was used for displaying these ED images. 4 × 4 binning was also applied for improving visualization of these ED images.

**Supplementary Figure 9.**
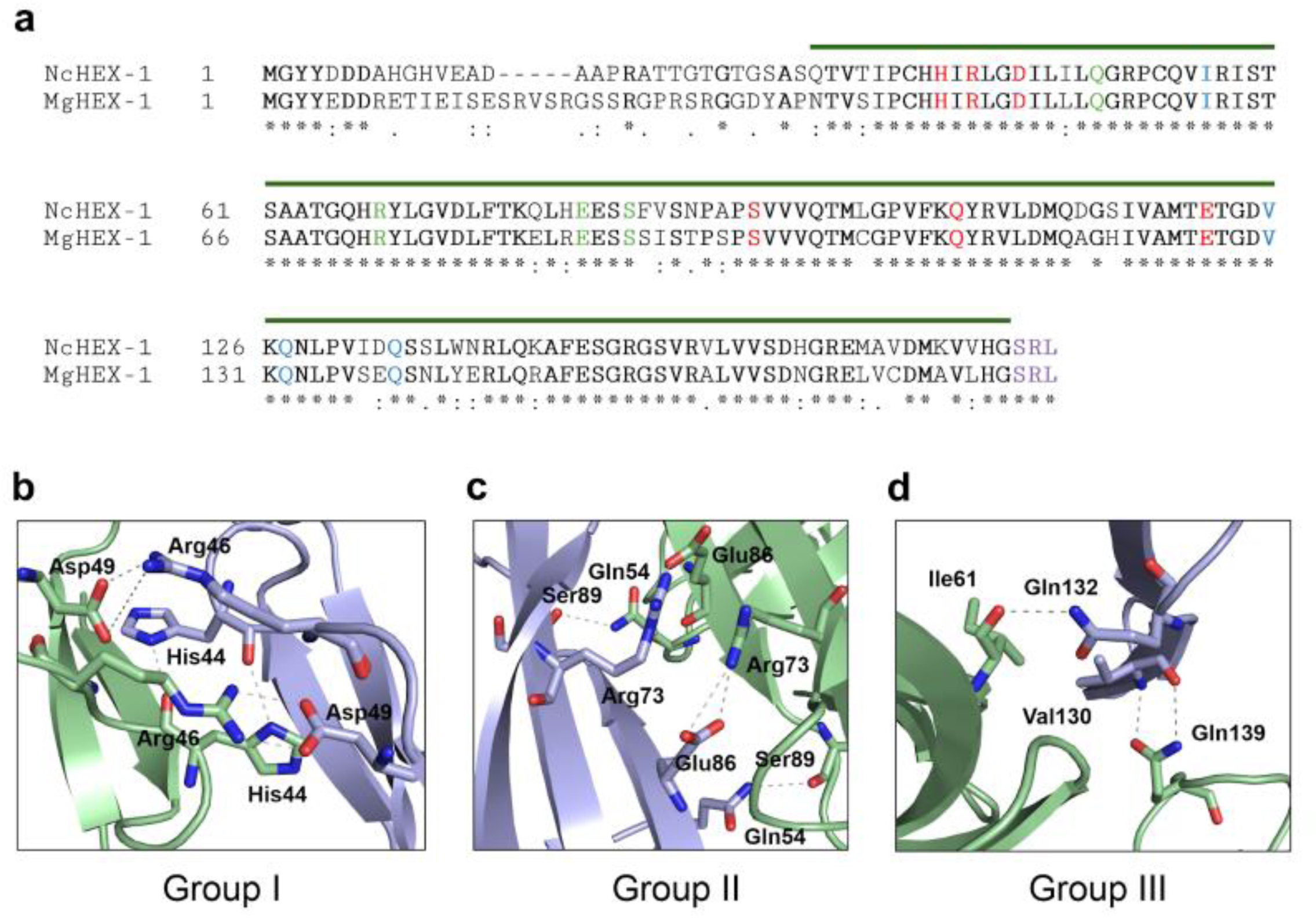
Conservation of *Mg*HEX-1 and *Nc*HEX-1 sequences and crystal lattice interactions. (**a**) Alignment of amino acid sequences of *Nc*HEX-1 with that of *Mg*HEX-1. Identical residues are marked by asterisks, similar residues by points. Residues that could be modelled in the electrostatic potential map calculated from the microvolume 3D ED data are indicated by the overlying green line. Residues involved in the stabilization of the crystal lattice of *Mg*HEX-1 are highlighted in red (Group I), green (Group II), and blue (Group III). The C-terminal SRL residues (violet) represent the peroxisome entry signal that was not resolved in the electron potential map. Details of the crystal lattice interactions are shown in (**b**) for Group I, (**c**) for Group II, and (**d**) for Group III interactions. The interacting monomers are shown in green and blue cartoon representation, while residues forming stabilizing hydrogen bonds (grey dashed lines) are highlighted as sticks.

## Supplementary Text 1

A comparison of *IncellED* and serial synchrotron-radiation X-ray diffraction (SSX) methods for structure elucidation of *in cellulo* MgHEX-1 crystals can be performed by taking into account the total volume of material used, the radiation dose absorbed and the multiplicity of the obtained diffraction data. Using these values from **Table 1** for the SSX and the microvolume 3D ED data, which yielded highly similar structural information, the efficiency advantage of 3D ED over SSX can be estimated as follows:

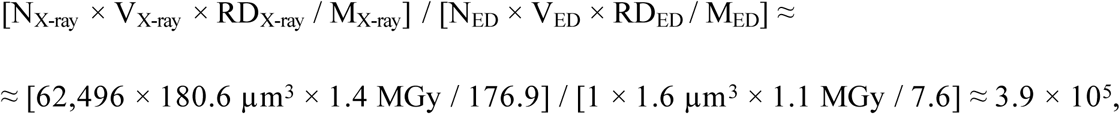

where N - number of single crystals used, V - volume of a single crystal used for data collection, RD - radiation dose absorbed, M - multiplicity of the diffraction datasets. The value for V_X-ray_ ≈ 7 × 3 × 8.6 µm^3^ ≈ 180.6 µm^3^ was evaluated considering the size of the X-ray beam (7 × 3 µm^2^; see *Methods*) and the average crystal size (∼8.6 µm).

